# Relative importance of MCL-1’s Anti-Apoptotic versus Non-Apoptotic Functions *in vivo*

**DOI:** 10.1101/2023.08.14.553217

**Authors:** Kerstin Brinkmann, Kate McArthur, Annli Tee, Andrew J. Kueh, Shezlie Malelang, Verena C. Wimmer, Leonie Gibson, Caitlin L Rowe, Philip Arandjelovic, Grant Dewson, Tracey L Putoczki, Philippe Bouillet, Naiyang Fu, Tim Thomas, Marco J. Herold, Anne K. Voss, Andreas Strasser

**Affiliations:** Blood Cells and Blood Cancer Division, The Walter and Eliza Hall Institute of Medical Research, Melbourne, 3052 Victoria, Australia; Department of Biochemistry and Molecular Biology, Biomedicine Discovery Institute, Monash University, Melbourne, 3800 Victoria, Australia; Epigenetics and Development Division, The Walter and Eliza Hall Institute of Medical Research, Melbourne, 3052 Victoria, Australia; ACRF Cancer Biology and Stem Cells Division, The Walter and Eliza Hall Institute of Medical Research, Melbourne, 3052 Victoria, Australia; Centre for Dynamic Imaging, The Walter and Eliza Hall Institute of Medical Research, Melbourne, 3052 Victoria, Australia; Department of Medical Biology, University of Melbourne, Melbourne, 3052 Victoria, Australia; Ubiquitin Signaling Division, The Walter and Eliza Hall Institute of Medical Research, Melbourne, 3052 Victoria, Australia; Personalised Oncology Division, The Walter and Eliza Hall Institute of Medical Research, Melbourne, 3052 Victoria, Australia; Department of Surgery, University of Melbourne, Melbourne, 3052 Victoria, Australia; Cancer and Stem Cell Biology Program, Duke-NUS Medical School, Singapore 169857, Singapore; Olivia Newton-John Cancer Research Institute, Heidelberg, Melbourne, 3084 Victoria, Australia; School of Cancer Medicine, La Trobe University, Bundoora, Melbourne, 3086 Victoria, Melbourne

## Abstract

MCL-1 is an anti-apoptotic member of the BCL-2 protein family that ensures cell survival by blocking the intrinsic apoptotic cell death pathway^1^. MCL-1 is unique in being essential for early embryonic development and the survival of many cell types, including many cancer cells, which are not affected by the loss of the other anti-apoptotic BCL-2 family members^1–4^. Non-apoptotic functions of MCL-1 controlling mitochondrial ATP production and dynamics have been proposed to underlie this unique requirement for MCL-1^5–9^. The relative contributions of the anti-apoptotic *versus* the non-apoptotic functions of MCL-1 in normal physiology have not been addressed. Here we replaced the coding sequence for MCL-1 with those for the anti-apoptotic proteins BCL-XL, BCL-2 or A1. We hypothesised that BCL-XL, BCL-2 and A1 may substitute for MCL-1 in the inhibition of apoptosis, but that they will not be able to replace MCL-1’s non-apoptotic function. Strikingly, *Mcl-1^Bcl-xL/Bcl-xL^* and *Mcl-1^Bcl-^*^2^*^/Bcl-^*^2^ embryos survived to embryonic day 14.5, greatly surpassing the pre-implantation lethality of *Mcl-1*^−/−^ embryos at E3.5. This demonstrates that the non-apoptotic functions of MCL-1 are dispensable for early development. However, at later stages of development and life after birth many cell types, particularly ones with high energy demand, were found to require both the anti-apoptotic and the non-apoptotic functions of MCL-1. These findings reveal the relative importance of these distinct functions of MCL-1 in physiology, providing important information for basic biology and the advancement of MCL-1 inhibitors in cancer therapy.

---

Apoptotic cell death is involved in morphogenesis of multi-cellular organisms^10,11^ and tissue homeostasis after birth^12^. This form of programmed cell death is utilised to remove superfluous, defunct and potentially dangerous cells^13^. Defects in apoptosis are implicated in the development of degenerative disorders, autoimmune disease and cancer^13,14^. Apoptosis is regulated by three structurally and functionally distinct subgroups of the BCL-2 protein family^15,16^. Anti-apoptotic BCL-2 family members (BCL-2, BCL-XL, MCL-1, A1, BCL-W) inhibit apoptosis by neutralizing the initiators of apoptosis, the so-called BH3 only proteins (e.g. tBID, BIM, PUMA, NOXA) and restraining the apoptosis effectors BAX and BAK^15,17,18^. These interactions prevent BAX and BAK from causing mitochondrial outer membrane permeabilization (MOMP) which unleashes the caspase cascade that coordinates the dismantling of the cell^19,20^.

Of the five anti-apoptotic proteins, MCL-1 has the most prominent function with its genetic removal or BH3 mimetic drug-mediated inhibition causing the death of many cell types, including early embryonic cells, that are not affected by the absence or inhibition of related anti-apoptotic BCL-2 proteins^2,4^. MCL-1-deficient embryos fail to implant and die around E3.5 ^9^, while BCL-XL-deficient embryos die around E13.5 due to aberrant death of erythroid progenitors and certain neuronal populations^21^. BCL-2-deficient mice die around 4 weeks after birth due to polycystic kidney disease caused by abnormal death of embryonic renal progenitors ^22,23^. A1 and BCL-W deficiency cause only minor defects^24^ (Extended Data Fig. 1a). It is not clear why mammals have multiple anti-apoptotic BCL-2 proteins. This could be to allow complex cell type and stress specific regulation of apoptosis. Differences between the various anti-apoptotic BCL-2 proteins include their binding affinity to the pro-apoptotic BCL-2 family members (Extended Data Fig. 1b), protein turnover (MCL-1 ∼30 min, A1 ∼10 min, BCL-2, BCL-XL, BCL-W >20 h) (Extended Data Fig. 1c) and subcellular localisation (Extended Data Fig. 1d)^25–27^. Some anti-apoptotic BCL-2 proteins reportedly also exert apoptosis-unrelated functions^28^. A proteolytically processed form of MCL-1^8,29–31^ and full-length BCL-XL^32–34^, but not BCL-2^35^, were shown to reside in the inter-mitochondrial membrane space where they can regulate mitochondrial dynamics and ATP production (Extended Data Fig. 1e). The physiological importance of the non-apoptotic roles of MCL-1 (and other BCL-2 proteins) are unclear, however, it was stated that the dying *Mcl-1^-/-^* blastocysts did not display hallmarks of apoptosis^9^. Failure of implantation was thus ascribed to the loss of the non-apoptotic functions of MCL-1 ^29^. Moreover, cardiomyocytes were reported to require both the anti-apoptotic and non-apoptotic function of MCL-1 to maintain heart function.

To advance MCL-1 inhibitors as cancer therapeutics, it is imperative to determine whether their reported on-target toxicities and efficacy against malignant cells are solely due to blocking their anti-apoptotic function or also due to impairing their non-apoptotic roles^3,36^. To explore functional redundancies and differences between MCL-1 and other anti-apoptotic BCL-2 proteins, we generated mice in which the *Mcl-1* gene was replaced by the coding sequences for BCL-XL, BCL-2 or A1 (referred to as gene-swap mice). Remarkably, BCL-XL and BCL-2, but not A1, could to a large extent replace MCL-1 during embryogenesis, suggesting that the essential role of MCL-1 during embryogenesis is to restrain apoptosis and that its reported non-apoptotic function is not required. However, a critical role of the non-apoptotic function of MCL-1 in facilitating optimal mitochondrial ATP production in cell populations with a high energy demand (e.g, hepatocytes) was found after birth.

## Gene-swap strategy to study MCL-1 function

The lethality of *Mcl-1^-/-^* mice around E3.5 offered an opportunity to examine the functional overlaps and distinctions between different pro-survival BCL-2 proteins in a physiological setting (Extended Data Fig. 1a). We hypothesised that MCL-1 can only be substituted by another BCL-2 family member if there is no unique essential function of MCL-1. To test this, we used a gene replacement strategy. Due to the complexities of the *Mcl-1*, *Bcl-2*, *Bcl-xL* and *A1* loci, it is not possible with current technology to maintain the exon/intron structure. Therefore, we used cDNA sequences encoding human BCL-2 (hBCL-2; termed *Mcl-1^Bcl-^*^2^ mice), FLAG-tagged mouse BCL-XL (*Mcl-1^Bcl-xL^*) or FLAG-tagged mouse A1 (*Mcl-1^A^*^1^) to replace the *Mcl-1* coding exons and intervening introns, all on a C57BL/6 background (Fig. 1a). Our strategy maintained the 5’ and 3’ untranslated regions of the *Mcl-1* gene locus. To verify that the loss of the *Mcl-1* introns did not have adverse impact, we used a cDNA encoding human MCL-1 (hMCL-1) to replace the mouse *Mcl-1* coding exons and introns to generate *Mcl-1^hMcl-^*^1^ control mice.

**Fig. 1.**
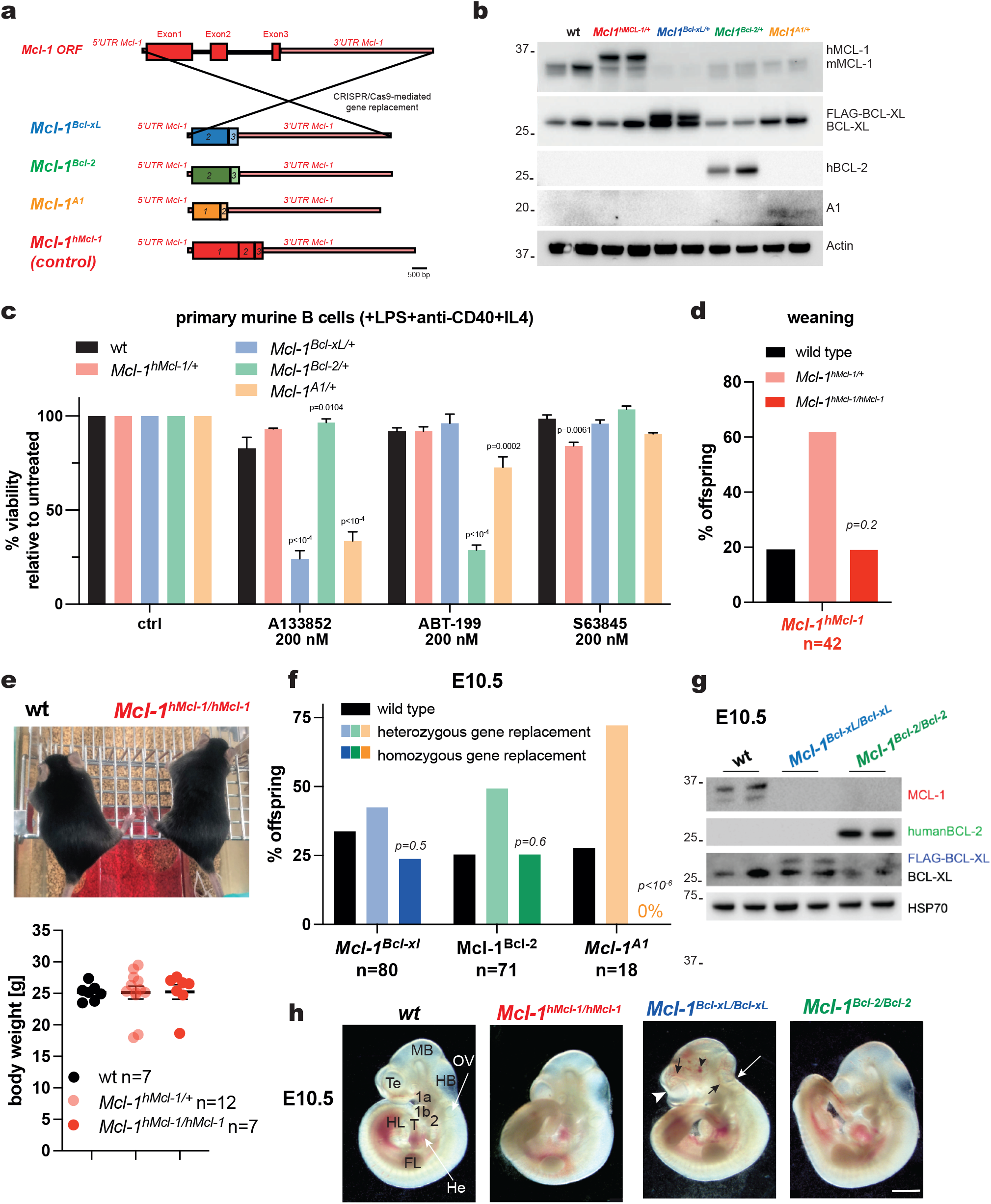
The essential function of MCL-1 in pre-implantation development can be replaced by BCL-XL or BCL-2 but not A1. **a**, Schematic presentation of the gene targeting strategy. **b**, Western blot of lysates from wild-type (wt), *Mcl-1^hMcl-^*^1^*^/+^*, *Mcl-1^Bcl-xL/+^*, *Mcl-1^Bcl-^*^2^*^/+^* and *Mcl-1^A^*^1^*^/+^* E10.5 embryos (n=2 per genotype, one embryo lysate per lane) detecting the indicated proteins. **c**, Survival of mitogen-activated B cells from *Mcl-1^hMcl-1/+^*, *Mcl-1^Bcl-xL/+^*, *Mcl-1^Bcl-2/+^* and *Mcl-1^A1/+^* mice treated with the indicated drugs for 24 h. Data are presented as mean ±SEM from 2 independent experiments, each performed in triplicates. Statistical significance was tested using two-way ANOVA with Tukey’s multiple comparison. p-values are indicated in the figure. **d**, Genotype frequencies at weaning among offspring of intercrosses of *Mcl-1^hMcl-1/+^* mice. **e**, Representative image showing a wild-type and Mcl-1hMcl-1/hMcl-1 mouse aged 50 days (top) and whole-body weight of 50-60 day-old wt (n=7), Mcl-1hMcl-1/+ (n=12) and Mcl-1hMcl-1/hMcl-1 (n=7) males (bottom). Statistical significance was tested using one-way ANOVA with Dunnet’s multiple comparison. **f**, Genotype frequencies at E10.5 among offspring of homotypic intercrosses of *Mcl-1^hMcl-1/+^*, *Mcl-1^Bcl-xL/+^*, *Mcl-1^Bcl-2/+^* or *Mcl-1^A1/+^* mice. **g**, Western blot of lysates from wild-type, *Mcl-1^hMcl-1/hMcl-1^*, *Mcl-1^Bcl-xL/Bcl-xL^* and *Mcl-1^Bcl-2/Bcl-2^* E10.5 embryos (n=2 per genotype, one embryo lysate per lane) detecting the indicated proteins. **h**, Representative images showing E10.5 wild-type, *Mcl-1^hMcl-1/hMcl-1^*, *Mcl-1^Bcl-xL/Bcl-xL^* and *Mcl-1^Bcl-2/Bcl-2^* embryos. **d**,**f**, Genotype distributions were analyzed computing the cumulative binomial probability of being less or equal to the expected value. p-values and number of animals (n) are indicated in the figure. **Labelling**: 1a, 1b, maxillary and mandibular portion of the 1st pharyngeal arch; 2 second pharyngeal arch; FL, forelimb bud; HB, hindbrain; He, heart, here beneath the tail; HL, hindlimb bud; MB, midbrain; OV, otic vesicle; T, tail; Te, telencephalon. White arrowhead and white arrow: areas of the lateral ventricles in the telencephalon and the 4th ventricle in the hindbrain, respectively, that appeared collapsed; black arrowhead: blood accumulation; black arrows: neuroepithelium forming ridges in the telencephalon and hindbrain.

## Validation of *Mcl-1* gene-swap strategy

Heterozygous *Mcl-1^hMcl^*^-1/+^*, Mcl-1^Bcl-x/+^, Mcl-1^Bcl-2/+^* and *Mcl-1^A1/+^* mice were viable and reached adulthood (Extended Data Fig. 2a). The expression of hMCL-1, FLAG-BCL-XL, hBCL-2 and FLAG-A1 protein was confirmed by Western blotting using whole E10.5 embryo lysates (Fig. 1b) and primary mouse embryonic fibroblasts (MEFs) (Extended Data Fig. 2c), as well as by flow cytometry using bone marrow cells (Extended Data Fig. 2b) from heterozygous gene-swap mice. We could discriminate between the endogenous mouse MCL-1 (mMCL-1) from the knock-in allele encoded hMCL-1 by differences in their molecular weight (hMCL-1 has a larger MW than mMCL-1^37^). The endogenous mouse BCL-2 (mBCL-2) protein could be discriminated from the knock-in allele encoded hBCL-2 by using monoclonal antibodies that specifically detect only mBCL-2 or only hBCL-2. The knock-in encoded FLAG-BCL-XL and FLAG-A1 could be discriminated from endogenous BCL-XL or A1 with an antibody detecting FLAG and their increased molecular weight seen when using antibodies against BCL-XL or A1.

The levels of endogenous BCL-2, BAX, BAK, PUMA and BIM remained unchanged in the E10 embryos, indicating that there were no compensatory mechanisms impacting these BCL-2 family members. However, we did observe a reduction in endogenous BCL-XL in *Mcl-1^Bcl-2/+^* E10.5 embryos lysates, and as expected, a reduction in MCL-1 in lysates from *Mcl-1^Bcl-x/+^, Mcl-1^Bcl-2/+^* E10.5 embryos and MEFs (Fig. 1b, Extended Data Fig. 2c-e).

To functionally validate the gene-swaps we treated mitogen-activated B cells with BH3 mimetic drugs (Fig. 1c). Activated B cells from *Mcl-1^hMcl-1/+^* mice were more susceptible to the MCL-1 inhibitor S63845 compared to cells from wild-type mice, consistent with the higher affinity of S63845 for hMCL-1 than mMCL-1^38^. Since B cells from heterozygous *Mcl-1^Bcl-xL/+^* or *Mcl-1^Bcl-2/+^* mice express higher total levels of BCL-XL (mBCL-XL + FLAG-BCL-XL) or BCL-2 (mBCL-2 + hBCL-2) and less MCL-1 compared to cells from wild-type mice, the former were more susceptible to the BCL-XL inhibitor A1331852 and the latter to the BCL-2 inhibitor ABT-199/Venetoclax. Collectively, these findings demonstrate the expression and functionality of the knock-in proteins in the gene-swap mice.

## Mcl-1^hMcl-1/hMcl-1^ mice are normal

To confirm that a gene-swap strategy at the *Mcl-1* gene locus is viable, we intercrossed *Mcl-1^hMcl-1/+^* mice and analysed their offspring. Homozygous *Mcl-1^hMcl-1/hMcl-1^* mice were born at a frequency that did not differ from the expected Mendelian frequencies (p=0.2; Fig. 1d) and these mice had normal appearance, weight and survival (Fig. 2e, Extended Data Fig. 3a). Histological analysis of the liver, kidney and heart revealed no abnormalities (Extended Data Fig. 3b-d). In accordance with previous analysis of humanised MCL-1 mice^37^, MEFs from *Mcl-1^hMcl-^*^1^*^/+^* and *Mcl-1^hMcl-1/hMcl-1^* mice showed normal responses to diverse apoptotic stimuli (Extended Data Fig. 3e). These findings demonstrate that the function of the mouse *Mcl-1* gene locus can be effectively replaced by a cDNA encoding human MCL-1, thereby validating our gene-swap strategy.

**Fig. 2.**
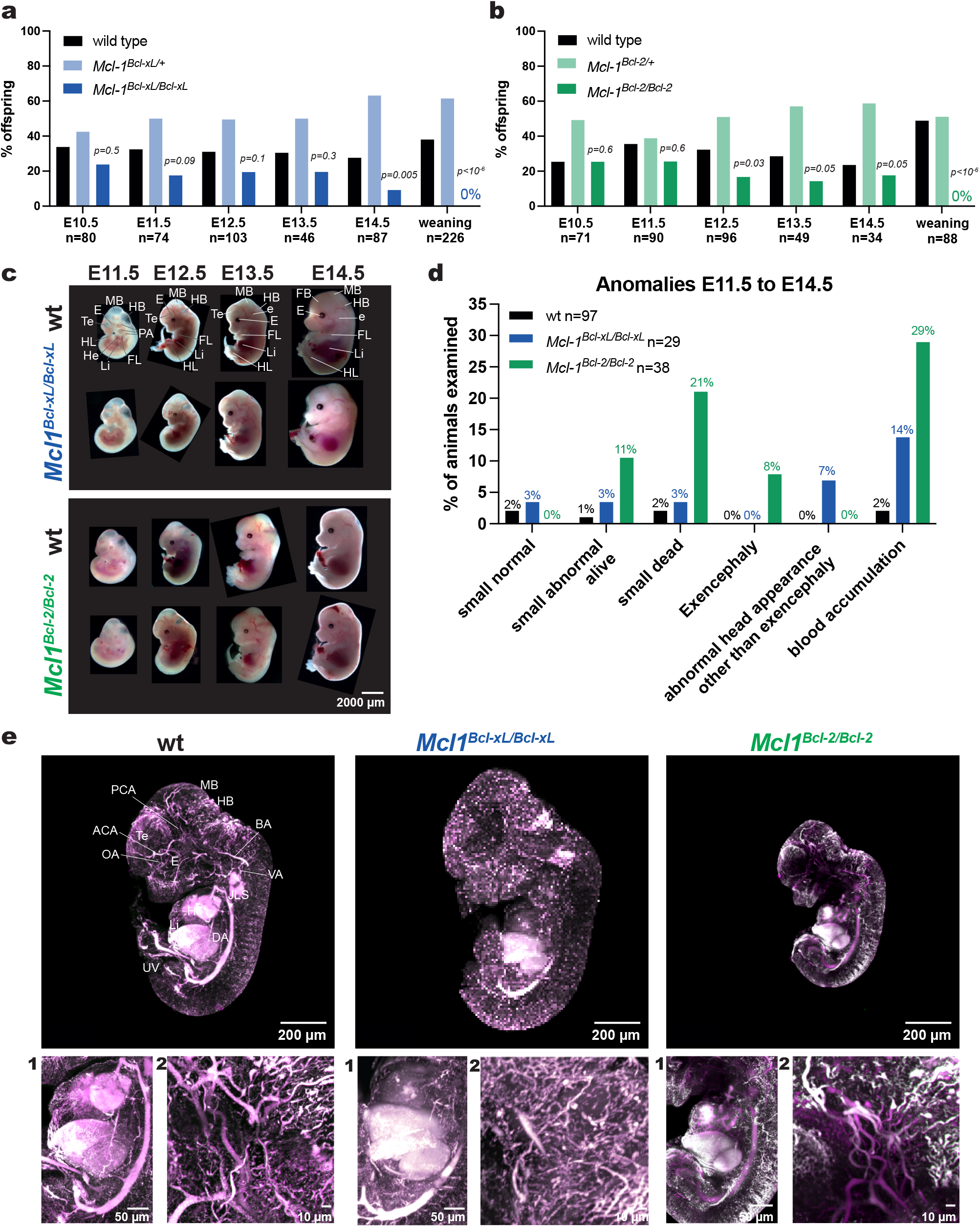
Homozygous *Mcl-1^Bcl-xL/Bcl-xL^*and *Mcl-1^Bcl-2/Bcl-2^* gene-swap embryos are more likely to exhibit abnormalities compared to wild-type embryos. **a, b,** Genotype frequencies at the indicated embryonic days among offspring of homotypic intercrosses of a, *Mcl-1^Bcl-xL/+^* or b, *Mcl-1^Bcl-2/+^* mice. Datasets for the offspring distribution at E10.5 are also presented in Fig. 1f and are included here for comparison. Genotype distributions were analysed computing the cumulative binomial probability of being less or equal to the expected value. p-values and number of animals (n) are indicated in the figure. **c**, Representative images showing wild-type, homozygous *Mcl-1^Bcl-xL/Bcl-xL^* and homozygous *Mcl-1^Bcl-2/Bcl-2^* embryos and foetuses at the indicated embryonic days. **d**, Summary frequencies of the phenotypes observed from E11.5 to E14.5 by external inspection of 29 *Mcl-1^Bcl-xL/Bcl-xL^*, 38 *Mcl-1^Bcl-2/Bcl-2^* and 97 wild-type embryos and foetuses at the indicated embryonic days. Data are presented as percentages of the total numbers of embryos per genotype. Numbers broken up by embryonic day are presented in Extended Data Table 1. **e,** Representative whole-mount images (iDISCO)^46^ showing a wild-type, a homozygous *Mcl-1^Bcl-xL/Bcl-xL^* and a homozygous *Mcl-1^Bcl-2/Bcl-2^*E12.5 embryo. The vasculature is marked by CD31/PCAM1 staining (purple) and blood cells appear white due to their inherent autofluorescence. Higher magnifications from each embryo are shown for (1) the developing heart and liver and (2) the developing cephalic vasculature. **Labelling**: ACA, anterior cerebral artery; BA, basilar artery; DA, descending aorta; E, eye; e, ear; FB, forebrain; FL, forelimb (forelimb bud, E11.5 and E12.5); HB, hindbrain; He, heart; HL, hindlimb (hindlimb bud, E11.5 and E12.5); JLS, jugular lymph sac, here blood filled; Li, liver; MCA, middle cerebral artery; MB, midbrain; OA, olfactory artery; PA, pharyngeal arches; PCA, posterior communicating artery; Te, telencephalon; UV, umbilical vein; VA, vertebral artery.

## Early embryogenesis without MCL-1

Next, we examined whether a rescue of the early embryonic lethality caused by the absence of MCL-1 could be achieved by replacing MCL-1 with hBCL-2, FLAG-BCL-XL or FLAG-A1. Intercrosses of heterozygous *Mcl-1^A1/+^* gene-swap mice did not yield any homozygous embryos at E10.5 or later stages of development (Fig. 1f, Extended Data Fig. 4a), possibly because the levels of FLAG-A1 were very low (Fig. 1b). In striking contrast, *Mcl-1^Bcl-xL/Bcl-xL^* and *Mcl-1^Bcl-2/Bcl-2^* homozygous gene-swap embryos were observed at E10.5 at frequencies that did not differ significantly from the expected Mendelian ratios (p=0.5 and 0.6, respectively; Fig. 2f). *Mcl-1^Bcl-2/Bcl-2^* or *Mcl-1^Bcl-xL/Bcl-xL^* embryos expressed hBCL-2 or FLAG-BCL-XL, respectively, but were devoid of MCL-1 (Fig. 1g). Hence, these embryos had successfully undergone the processes of pre-implantation development, implantation, gastrulation and embryonic patterning without any MCL-1 protein. The presence of the forebrain, midbrain, hindbrain and spinal cord in *Mcl-1^Bcl-xL/Bcl-xL^* and *Mcl-1^Bcl-2/Bcl-2^* embryos at E10.5 indicate that they had undergone neurulation and patterning of the neural tube (Fig. 1h). The normal size of the E10.5 embryos compared to controls and the observed heartbeat indicate that heart development and placentation were adequate to support development to this stage. However, anomalies were observed in some *Mcl-1^Bcl-xL/Bcl-xL^* and *Mcl-1^Bcl-2/Bcl-2^* embryos in the head region (collapsed lateral and 4th ventricles, ridged neuroepithelium; Fig. 1h). These findings demonstrate that BCL-2 and BCL-XL can replace MCL-1 during the essential pre-implantation state when 100% of *Mcl-1^-/-^* embryos die. In contrast to MCL-1 and BCL-XL, BCL-2 does not localise to the mitochondrial inter-membrane space^35^. Therefore, the presence of *Mcl-1^Bcl-2/Bcl-2^* embryos at E10.5 demonstrates that the reported non-apoptotic functions of MCL-1^29^ are not critical for early embryonic development.

## _Mcl-1Bcl-2/Bcl-2 vs Mcl-1Bcl-xL/Bcl-xL mice_

Homozygous *Mcl-1^Bcl-xL/Bcl-xL^* foetuses were under-represented at E14.5 (p=0.005) (Fig. 2a) and *Mcl-1^Bcl-2/Bcl-2^* embryos were already moderately under-represented from E12.5 onward (p=0.003 to 0.05; Fig. 2b). At E15.5 no live *Mcl-1^Bcl-xL/Bcl-xL^* or *Mcl-1^Bcl-2/Bcl-2^* foetuses were recovered, and *Mcl-1^Bcl-xL/Bcl-xL^* and *Mcl-1^Bcl-2/Bcl-2^* mice were absent at weaning (both p<10^-6^; Fig. 2a, b).

From E11.5 to E14.5 significantly higher percentages of abnormal *Mcl-1^Bcl-xL/Bcl-xL^* and *Mcl-1^Bcl-2/Bcl-2^* embryos and foetuses were observed compared to wild-type littermates (24% p=0.002 and 53% p<10^-6^ for *Mcl-1^Bcl-xL/Bcl-xL^* and *Mcl-1^Bcl-2/Bcl-2^*, respectively; 6% in wild-type controls; Fig. 2d, Extended Data Fig. 4b-d, Extended Data Table 1). Some homozygous gene-swap embryos were small and had an abnormal appearance (3% and 11% for *Mcl-1^Bcl-xL/Bcl-xL^* and *Mcl-1^Bcl-2/Bcl-2^*, respectively) and some were dead (3% and 21%; Fig. 2d, Extended Data Fig. 5, Extended Data Table 1). Anomalies observed included exencephaly (0% and 8%) and other abnormalities in the head region (7% and 0%; collapsed lateral and 4th ventricles, ridged neuroepithelium). Blood accumulation consistent with haemorrhages, mostly in the head region, was observed in 14% of *Mcl-1^Bcl-xL/Bcl-xL^* and 29% of *Mcl-1^Bcl-2/Bcl-2^* embryos and foetuses (Fig. 2d, Extended Data Fig. 4b-d, Extended Data Table 1).

Whole mount 3D imaging (iDISCO)^39^ of E12.5 embryos (Fig. 2e; Extended Data Movies 1-3) upon staining of endothelial cells together with the inherent auto-fluorescence of blood revealed no major differences in vital organs, such as the heart or foetal liver, and vascularisation (Fig. 2e, zoom in 1-2) between genotypes in the specimens examined.

Notably, a considerably smaller proportion of normal *Mcl-1^Bcl-2/Bcl-2^* embryos was observed compared to *Mcl-1^Bcl-xL/Bcl-xL^* embryos (p=0.0001), and exencephaly was only observed in *Mcl-1^Bcl-2/Bcl-2^* embryos (Fig. 2d, Extended Data Fig. 4b-d, Extended Data Table 1). This indicates that at later stages of development a non-apoptotic function of MCL-1 is critical; and BCL-XL may (at least partially) replace the reported function of MCL-1 in mitochondrial energy production and dynamics, whereas BCL-2 is not able to rescue this non-apoptotic function.

## *Mcl-1* gene-swap reduces apoptosis

The defects in homozygous *Mcl-1^Bcl-xL/Bcl-xL^* and *Mcl-1^Bcl-2/Bcl-2^* embryos at later developmental stages (Fig. 2) could be due to impaired apoptosis since a short-lived anti-apoptotic protein (MCL-1) was replaced with a presumably more potent long-lived relative (BCL-2, BCL-XL). Importantly, loss of the pro-apoptotic effectors BAX, BAK and (BOK) causes severe and often lethal developmental defects^40,41^. Accordingly, MEFs and mitogen activated B cells derived from *Mcl-1^Bcl-xL/+^* or *Mcl-1^Bcl-2/+^* embryos and mice, respectively, were significantly more resistant to etoposide- and cytarabine-induced cell death compared to their wild-type counterparts (Fig. 3a, Extended Data Fig. 5a). Strikingly, heterozygous *Mcl-1^Bcl-xL/+^* and *Mcl-1^Bcl-2/+^* females displayed an abnormally high incidence of vaginal septa (p<10^-4^ for both), an indicator of abnormally reduced apoptosis during development (Fig. 3b).

**Fig. 3.**
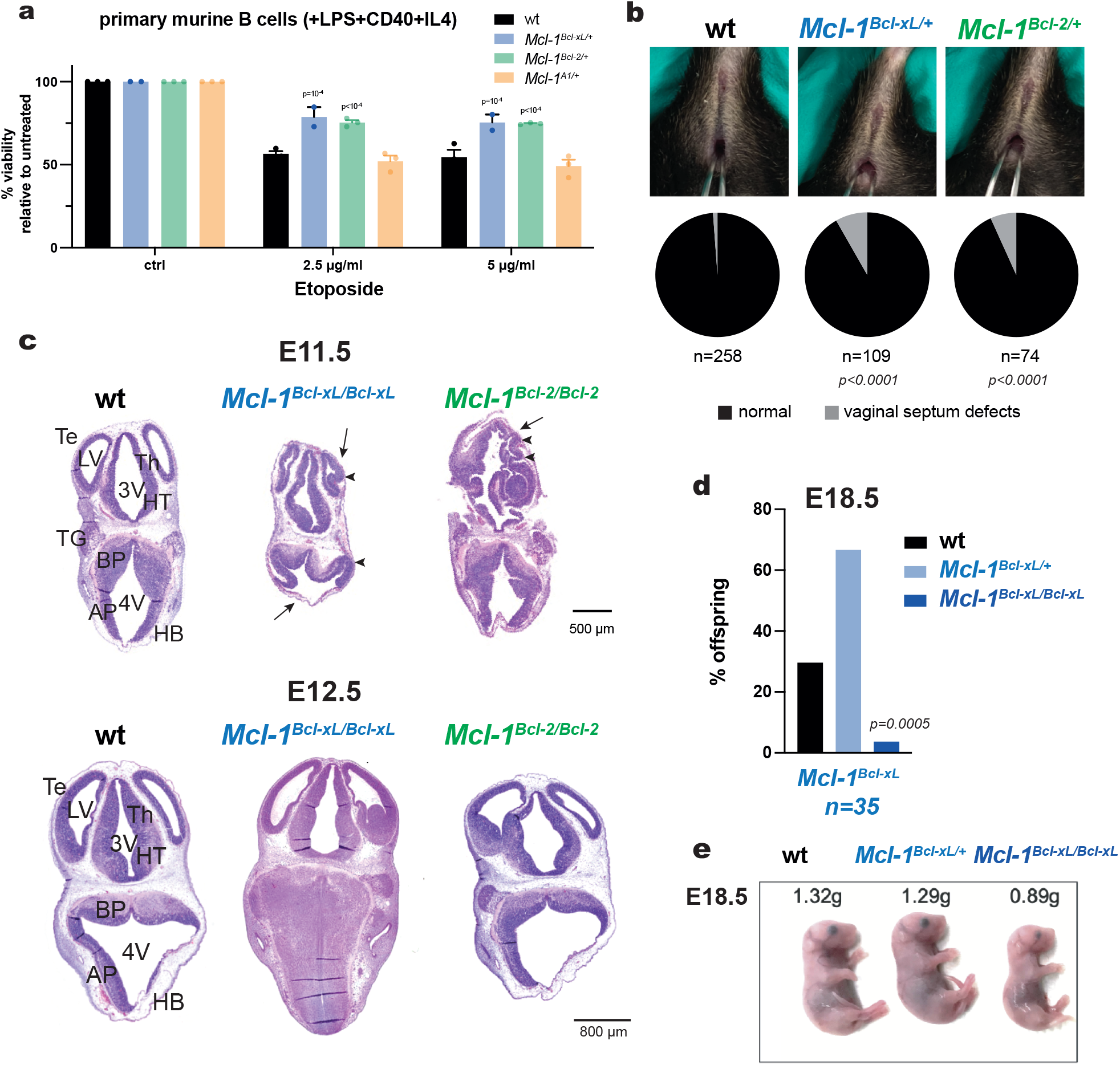
Cells from *Mcl-1^Bcl-xL^*and *Mcl-1^Bcl-2^* gene-swap mice show increased resistance to apoptosis. **a,** Survival analysis of mitogen activated B cells from wild-type (wt, n=3), heterozygous *Mcl-1^Bcl-xL/+^*(n=2), *Mcl-1^Bcl-2/+^* (n=3) and *Mcl-1^A1/+^* (n=3) gene-swap mice. Mitogen activated B cells were treated with the indicated concentrations of cytarabine and cell viability was assessed after 24 h by flow cytometry after staining with annexin V and propidium iodide. Data are presented as mean ± SEM from ≥ 2 independent experiments, each performed in triplicates. Statistical significance was tested using two-way ANOVA with Tukey’s multiple comparison. p-values are indicated in the figure. **b**, Representative images showing the vaginal septa of a female wild-type, heterozygous *Mcl-1^Bcl-xL/+^* and *Mcl-1^Bcl-2/+^*mouse (top row) and incidence of vaginal septa defects in wild-type (n=258), heterozygous *Mcl-1^Bcl-xL/+^* (n=109) and heterozygous *Mcl-1^Bcl-2/+^*(n=74) females (bottom row). Statistical significance was tested using the Chi-square test comparing the observed frequencies and the expected frequencies (wild-type). p-values are indicated in the figure. **c**, Representative images of H&E-stained serial sections of homozygous *Mcl-1^Bcl-xL/Bcl-xL^* and homozygous *Mcl-1^Bcl-2/Bcl-2^* embryos at embryonic days E11.5 and E12.5 at the forebrain/hindbrain level. Images of wild-type littermates are shown for comparison. **d**, Genotype frequency at E18.5 among offspring of intercrosses of heterozygous *Mcl-1^Bcl-xL/+^* mice. Genotype distributions were analysed computing the cumulative binomial probability of being less or equal to the expected value. p-values and number of foetuses (n) are indicated in the figure. **e**, Images showing a wild-type, a heterozygous *Mcl-1^Bcl-xL/+^* and a homozygous *Mcl-1^Bcl-xL/Bcl-xL^* embryo at embryonic day E18.5 with whole-body weights indicated. **Labelling**: 3V, third ventricle; 4V, mesencephalic vesicle (future 4th ventricle); AP and BP, alar and basal plate of the hindbrain; HB, hindbrain; HT, hypothalamus (diencephalon); LV, telencephalic vesicle (future lateral ventricle); Th, thalamus (diencephalon); Te, telencephalon.

Histological analysis of serial sections revealed that some E11.5, E12.5 and E13.5 *Mcl-1^Bcl-xL/Bcl-xL^* and *Mcl-1^Bcl-2/Bcl-2^* embryos displayed folding of the neuroepithelium in several areas of the developing brain (Fig. 3C, Extended Data Fig. 5b), which appeared to be due to the collapse of the lateral and 4th brain ventricles. At E14.5, we did not observe folding of the neuroepithelium in *Mcl-1^Bcl-2/Bcl-2^* foetuses (Extended Data Fig. 5c). We did not detect any defects in neuronal tube closure in the *Mcl-1^Bcl-xL/Bcl-xL^* embryos assessed, whereas 8% of *Mcl-1^Bcl-2/Bcl-2^* embryos presented with exencephaly (Fig. 2d, Extended Data Fig. 4b-d), a phenotype that is also observed in mice with a complete block in apoptosis^41^. These findings indicate that replacement of MCL-1 with BCL-2, and to a lesser extent with BCL-XL can cause developmental anomalies through abnormally increased cell survival, possibly due to longer protein lifespan and differences in interactions with pro-apoptotic BCL-2 family members. However, of note some *Mcl-1^Bcl-xL/Bcl-xL^* and *Mcl-1^Bcl-2/Bcl-2^* foetuses were lost before E15.5 and none reach weaning (Fig. 2a, b), whereas many *Bax^-/-^Bak^-/-^Bok^-/-^* triple knock-out (TKO) embryos lacking all executioners of intrinsic apoptosis develop until E18.5 and 1.8% even reach weaning (Extended Data Fig. 5d)^41^. Thus, while some abnormalities might be attributed to apoptotic defects during development, the failure of *Mcl-1^Bcl-xL/Bcl-xL^* and *Mcl-1^Bcl-2/Bcl-2^* to survive beyond E15.5, unlike the *Bax^-/-^Bak^-/-^Bok^-/-^* TKO mice, may be caused by the absence of non-apoptotic functions of MCL-1.

## Mixed background *Mcl-1^Bcl-xL/Bcl-xL^* pups

One homozygous *Mcl-1^Bcl-xL/Bcl-xL^* pup developed to E18.5 (Fig. 3d). Although it was smaller than its wild-type and heterozygous littermates, it looked otherwise remarkably normal (Fig. 3e). Since a mixed genetic background can be more tolerant of genetic abnormalities and allow further development (e.g. *Phf6*^42,43^ or *Tgfb1* mice^44^) we bred *Mcl-1^Bcl-2^* and *Mcl-1^Bcl-xL^* gene-swap mice on a mixed FVBxBALB/cxC57BL/6 genetic background.

Importantly, the absence of MCL-1 also causes pre-implantation lethality on this mixed background (Extended Data Fig. 6a, b). Although underrepresented (p=0.04), 18% *Mcl-1^-/-^* embryos were observed at E2.5 (Extended Data Fig. 6a, b). However, at this stage *Mcl-1^-/-^* embryos were already compromised and rarely reached the morula stage. *Mcl-1^-/-^* embryos failed to develop to the E3.5 blastocysts stage (p<10^-6^) and did not produce inner cell mass outgrowths in culture (p<10^-6^; Extended Data Fig. 6a, b). In contrast to the initial report^9^, we detected TUNEL^+^ apoptotic cells in a dying *Mcl-1^-/-^* E2.5 morula (Extended Data Fig. 6c-e), indicating that the lethality is presumably caused by aberrant apoptotic cell death.

Remarkably, many homozygous *Mcl-1^Bcl-xL/Bcl-xL^* mice were born alive on the mixed genetic background and, although underrepresented (p=4x10^-6^), 11% reached weaning (Fig. 4a). Interestingly, no homozygous *Mcl-1^Bcl-2/Bcl-2^* mice were born alive (p<10^-6^) or present at E19.5 (p=0.01) (Fig. 4b). This finding indicates that the reported non-apoptotic functions of MCL-1 are critical for the survival of mice during the foetal phase of development and after birth and that this function can be replaced, albeit imperfectly, by BCL-XL but not by BCL-2.

**Fig. 4.**
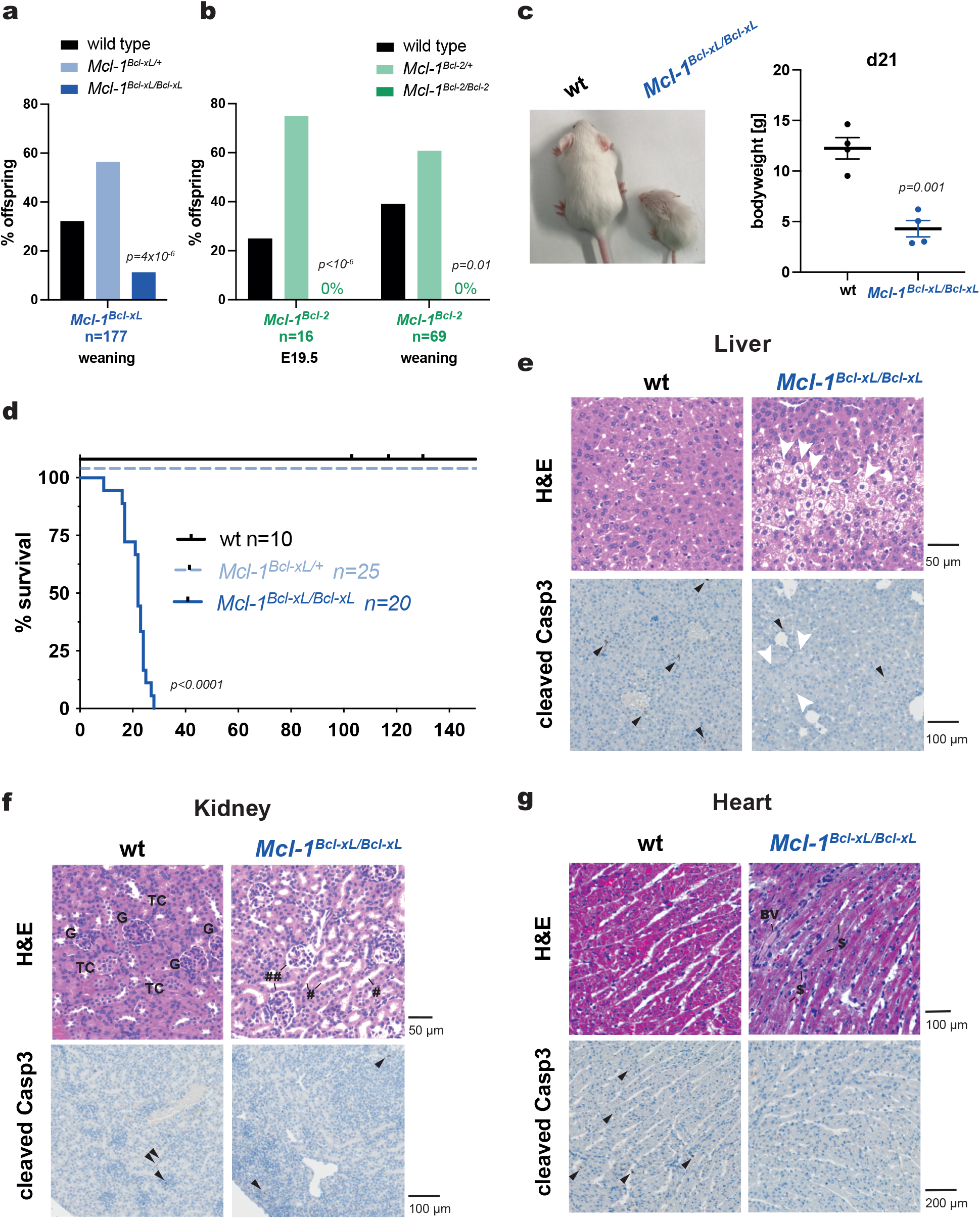
Loss of the non-apoptotic function of MCL-1 is critical for survival of mice after birth. **a,** Genotype frequency at weaning among offspring of intercrosses between *Mcl-1^Bcl-xL/+^* (n=177) mice (FVBxBALB/cxC57BL/6 mixed genetic background). Genotype distributions were analysed as described above. **b**, Genotype frequency among offspring of intercrosses between *Mcl-1^Bcl-2/+^*mice (FVBxBALB/cxC57BL/6 mixed genetic background) at weaning (n=69) and E19.5 (n=16). Genotype distributions were analysed as described above. p-values are indicated in the figure. **c**, Representative image showing a wild-type and a *Mcl-1^Bcl-xL/Bcl-xL^*mouse at day 21 after birth (FVBxBALB/cxC57 BL/6 mixed genetic background, left panel). Body weights of wild-type (n=4) and *Mcl-1^Bcl-xL/Bcl-xL^* (n=4) mice were assessed at day 21 after birth (FVBxBALB/cxC57BL /6 mixed genetic background, male and female, right panel). Statistical significance was tested using a two-tailed t-test. p-values are indicated in the figure. **d**, Kaplan-Meyer survival curve showing the survival of wild-type (n=10), heterozygous *Mcl-1^Bcl-xL/+^*(n=25) and homozygous *Mcl-1^Bcl-xL/Bcl-xL^* (n=20) mice (FVBxBALB/cxC57 BL/6 mixed genetic background). Statistical significance was tested using the Log Rank (Mantle-Cox) test. p-values are indicated in the figure. **e-g**, Representative histological images of H&E-stained sections (top row) and sections stained for cleaved (activated) caspase3 (bottom row) of wild-type (left panel) and homozygous *Mcl-1^Bcl-xL/Bcl-xL^* mice (right panel) (FVBxBALB/cxC57BL/6 mixed genetic background) of **e**, liver, **f**, kidney and **g**, heart myocardium. Black arrowheads indicate apoptotic (cleaved caspase-3 positive) cells. White arrowheads indicate ballooning hepatocytes. # indicate enlarged renal tubule lumina enclosed by degenerating renal tubular cells. ## indicate degenerating glomeruli. $ indicate degenerating cardiomyocytes. **Labelling**: BV, blood vessel; G, glomeruli; GC, goblet cell; TC, tubular cells.

## BCL-XL cannot fully replace MCL-1 in pups

Homozygous *Mcl-1^Bcl-xL/Bcl-xL^* pups were abnormally small, displayed a hunched posture and an enlarged head but looked otherwise normal (Fig. 4c). All *Mcl-1^Bcl-xL/Bcl-xL^* pups died within 30 days from birth (Fig. 4d). Histological examination failed to reveal marked structural abnormalities in the brains of the runted *Mcl-1^Bcl-xL/Bcl-xL^* pups (Extended Data Fig. 7a). In contrast, other vital organs, including liver (Fig 4e), kidneys (Fig 4f), heart (Fig 4g) and intestines (Extended Data Fig. 7e), showed substantial defects. The most dramatic defects were observed in the livers of the runted *Mcl-1^Bcl-xL/Bcl-xL^* pups. They contained large numbers of ballooning hepatocytes (Fig. 4e), which may indicate metabolic defects^45^. In the small intestine, disorganised and cell-poor crypts were observed (Extended Data Fig. 7e). The kidney damage appeared to mainly comprise degeneration of proximal renal tubular cells and glomeruli (Fig. 4f), a cell type displaying high metabolic activity^46^. In the heart, cardiomyocytes appeared to undergo degeneration (Fig. 4g). Staining for activated (cleaved) caspase-3 (Fig 4e-g and Extended Data Fig. 7e, bottom rows) and TUNEL immunohistochemistry of the liver (Extended Data Fig. 7d) revealed no increased apoptosis in the impacted organs, suggesting that the *Mcl-1^Bcl-xL/Bcl-xL^* defects are caused by the loss of a non-apoptotic function of MCL-1. No severe cardiac fibrosis was observed (Extended Data Fig. 7b), a phenotype that was observed upon inducible whole-body deletion of MCL-1 in *Mcl-1^fl/fl^RosaCreERT2* mice (Extended Data Fig. 7c). This is in agreement with the cardiomyocyte specific deletion of MCL-1 (*Mcl-^fl/fl^Ckmm-Cre*)^8^ where the fibrotic phenotype was shown to be driven by aberrant apoptosis^8^.

## MCL-1 is critical for liver homeostasis

Based on the relative severity of the anomalies in the different tissues, we focussed on the liver for a more detailed analysis. First, we confirmed that no MCL-1 protein was present in the livers of *Mcl-1^Bcl-xL/Bcl-xL^* pups by immunofluorescence staining (Extended Data Fig. 7f). Intriguingly, the phenotype of the runted *Mcl-1^Bcl-xL/Bcl-xL^* pups was similar to *Vdac2^-/-^* pups which also died at a young age and similarly presented with ballooning hepatocytes and liver damage (Fig. 5a)^47^. VDAC2 is a voltage dependent ion channel localised in the mitochondrial outer membrane where it facilitates the shuttling of mitochondrial produced ATP into the cytosol^48^. Given the similarities of the runted *Mcl-1^Bcl-xL/Bcl-xL^* and *Vdac2^-/-^* pups, we hypothesised that BCL-XL may not be sufficient to fully replace MCL-1’s proposed role in mitochondrial ATP production^7,8,29^. To explore this, we compared the metabolic capacity of wild-type, *Mcl-1^-/-^* ^49^ and *Mcl-1^Bcl-xL/Bcl-xL^* MEFs. We confirmed that *Mcl-1^Bcl-xL/Bcl-xL^* MEFs express FLAG-BCL-XL but no MCL-1 protein (Fig. 5b) and, as predicted, are highly sensitive to the BCL-XL inhibitor A1131852 (Extended Data Fig. 8a). In line with previous reports^29^, *Mcl-1^-/-^* MEFs had defects specifically in complex IV assembly when compared to wild-type MEFs. Remarkably, this defect was rescued in *Mcl-1^Bcl-xL/Bcl-xL^* MEFs (Fig. 5c, Extended Data Fig. 8b, c). Furthermore, unlike *Mcl-1^-/-^* MEFs which display defective cristae and balloon-like structures at the ultrastructural level^7,8,29^, mitochondria from *Mcl-1^Bcl-xL/Bcl-xL^* MEFs showed normal cristae organisation (Fig. 5d). Strikingly, we also observed no significant changes in complex IV assembly in liver samples from the runted *Mcl-1^Bcl-xL/Bcl-xL^*pups when compared to their wild-type littermates (Fig. 5e). This demonstrates that at least in MEFs and hepatocytes BCL-XL is able to substitute for the reported function of MCL-1 in the assembly of the mitochondrial electron transport chain. Nevertheless, *Mcl-1^Bcl-xL/Bcl-xL^* pups presented with severe liver damage, that is likely the result of metabolic defects (Fig. 4e, 5a). Indeed, an additional unique non-apoptotic function of MCL-1 in the regulation of fatty acid oxidation (FAO) has been reported^6^. In line with this study, OilRedO-staining of liver sections from *Mcl-1^Bcl-xL/Bcl-xL^* pups revealed a dramatic accumulation of lipids in hepatocytes (Fig. 5f) that may result from the insufficient fatty acid break down due to the reported downregulation of the FAO pathway caused by the absence of MCL-1^6^.

**Fig. 5.**
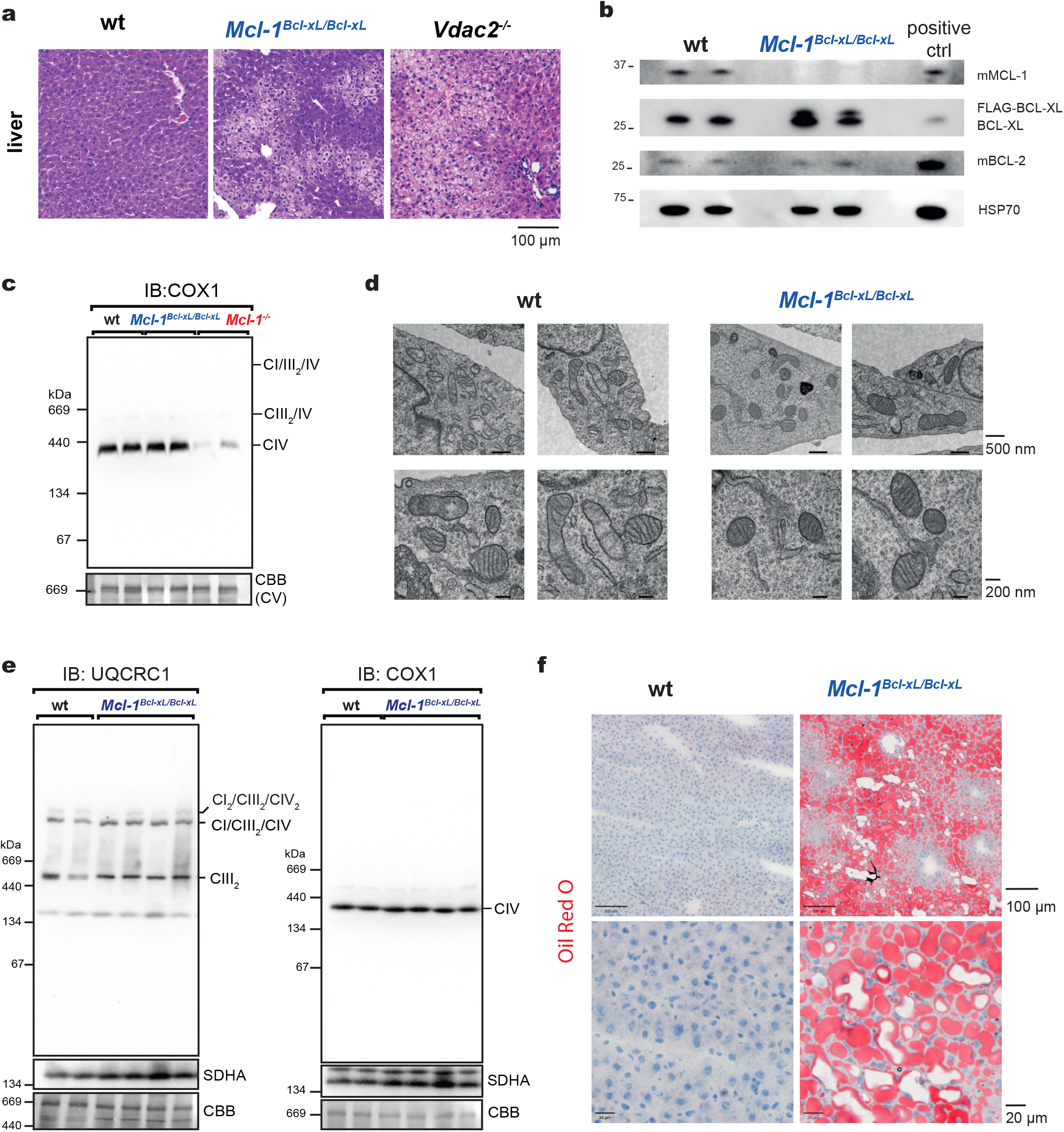
MCL-1’s non-apoptotic function is critical for liver homeostasis. **a**, Representative H&E-stained liver sections from wild-type (left panel), homozygous *Mcl-1^Bcl-xL/Bcl-xL^* (middle panel) and *Vdac2^-/-^*mice (right panel). **b**, Western blot analysis of wild-type and homozygous *Mcl-1^Bcl-xL/Bcl-xL^* MEFs (n=2 per genotype) detecting mouse MCL-1), BCL-XL (detecting both endogenous mouse BCL-XL and FLAG-BCL-XL encoded from the *Mcl-1* locus; note, the latter runs at a higher molecular weight), mouse BCL-2 and HSP70 (loading control). MEFs were immortalised using the SV40 large T antigen. The murine *Eµ-Myc* double-hit lymphoma (DHL) cell line 214DHL (known to express high levels of mBCL-2 and moderate levels of mMCL-1) was used as positive control. **c**, Blue native PAGE and immunoblot analysis for respiratory complex IV (CIV) (using an antibody to COX1) on isolated mitochondria from wild-type, *Mcl-1^-/-^*and *Mcl-1^Bcl-xL/Bcl-xL^* MEFs (n=2 each genotype). A portion of the Coomassie stained blot (CBB) with complex V (CV) shown serves as a loading control. A long exposure of the same blot is presented in Extended Data Fig. 9b. **d**, Representative transmission electron microscopy (TEM) images from wild-type and *Mcl-1^Bcl-xL/Bcl-xL^* MEFs, at either 10,000x or 20,000x magnification (as indicated). **e**, Blue native PAGE and immunoblot analysis for respiratory complex I, II, III (CI, CII, CIII) (using an antibody to UQCRC1) and complex IV (CIV) (using an antibody to COX1), on isolated mitochondria from wild-type and *Mcl-1^Bcl-xL/Bcl-xL^* hepatocytes. A portion of the Coomassie stained blot (CBB) with complex V (CV) shown serves as a loading control. **f**, Representative image of OilRedO stained liver sections (detecting lipids) from wild-type (left panel) and homozygous *Mcl-1^Bcl-xL/Bcl-xL^* pups (right panel).

These findings demonstrate that in several tissues of newborn mice MCL-1 is critical for both, inhibiting apoptosis as well as regulating a vital non-apoptotic process. Furthermore, our data demonstrate that in contrast to BCL-2, BCL-XL can substitute for MCL-1 in the regulation for the mitochondrial electron transport chain. However, neither BCL-XL nor BCL-2 can substitute for MCL-1 in the regulation of the FAO pathway.

## Discussion

Our gene-swap strategy resolved the physiological importance of the anti-apoptotic *versus* non-apoptotic functions of MCL-1. Notably, BCL-XL has also been reported to regulate mitochondrial ATP production^32,33^, whereas BCL-2 is not expected to have such a role because it is not localised in the inter-mitochondrial membrane space^35,50^. Therefore, our gene-swap mice made it possible to discriminate the effects caused by loss of the anti-apoptotic function of MCL-1 *versus* its role in mitochondrial ATP-production.

Our findings provide evidence that both the anti-apoptotic and the non-apoptotic functions of MCL-1 are critical for foetal development and life after birth. Interestingly, the relative importance of these functions differs across developmental stages and cell types. The failure of *Mcl-1^-/-^* blastocysts to implant was reported to occur in the absence of apoptosis^9^. However, the replacement of MCL-1 with either BCL-2 or BCL-XL rescued embryos until E13.5. This indicates that only the anti-apoptotic but not the non-apoptotic function of MCL-1 is essential for early embryogenesis. Congruently, we found evidence of apoptotic DNA fragmentation in a *Mcl-1^-/-^* E2.5 embryo. These findings indicate that the machinery to undergo apoptosis is active at E2.5 and thus *Mcl-1^-/-^* embryos die of aberrant apoptosis.

*Mcl-1^Bcl-xL/Bcl-xL^* and *Mcl-1^Bcl-^*^2^*^/Bcl-^*^2^ foetuses died significantly earlier than foetuses lacking all executioners of apoptosis (*Bax^-/-^Bak^-/-^Bok^-/-^*)^41^. This indicates that abnormalities causing the death of *Mcl-1^Bcl-xL/Bcl-xL^* and *Mcl-1^Bcl-2/Bcl-2^* foetuses cannot solely result from increased cell survival due to the replacement of short-lived MCL-1 with long-lived BCL-2 or BCL-XL. Our findings thus support a critical non-apoptotic role of MCL-1 *in vivo*. On a mixed genetic background *Mcl-1^Bcl-xL/Bcl-xL^* but no *Mcl-1^Bcl-^*^2^*^/Bcl-2^* mice were born. We propose that these differences are due to the ability of BCL-XL, but not BCL-2, to boost mitochondrial ATP production ^32,33^. Accordingly, on a C57BL/6 background, a significantly smaller proportion of normal *Mcl-1^Bcl-2/Bcl-2^* embryos at E14.5 and E15.5 was observed compared to *Mcl-1^Bcl-xL/Bcl-xL^* mice.

In line with some previous reports^32,33^, we observed that BCL-XL can substitute for MCL-1 in the assembly of the mitochondrial electron transport chain. Nevertheless *Mcl-1^Bcl-xL/Bcl-xL^* pups on the mixed genetic background presented with fatal liver pathology characterised by ballooning hepatocytes and lipid accumulation in the absence of aberrant hepatocyte apoptosis. This indicates that MCL-1 may have an additional function in the regulation of metabolic processes that cannot be rescued by BCL-XL. Notably, such an additional function of MCL-1 in the regulation of the fatty acid oxidation (FAO) pathway was recently described^6^. Specifically, MCL-1 was reported to enforce a dependency on FAO in leukaemic cells and its loss resulted in downregulation of enzymes involved in FAO^6^.

Overall, our findings suggest that differences in gene expression patterns, protein half-life and non-apoptotic functions may provide the appropriate control of the survival of diverse cell types at distinct stages of differentiation when they are subject to different conditions of stress and energy demand. Finally, our findings lend support for the idea^51^ that the BCL-2 protein family may originally have evolved to increase energy production in eukaryotic cells when they first developed mitochondria, and that the functions of BCL-2 family proteins in the control of apoptosis only emerged subsequently. Certain BCL-2 family members, including MCL-1 and BCL-XL, appear to have retained some of this original function^51,52^.

## Methods

### Mice

*Mcl-1^hMcl-1^, Mcl-1^Bcl-xL^, Mcl-1^Bcl-2^ and Mcl-1^A1^* mice were generated on a C57BL/6 J background by the MAGEC laboratory at The Walter and Eliza Hall Institute, as previously described^53^. 20 ng/μL Cas9 mRNA, 10 ng/μL single guide RNAs and 40 ng/μL donor template were injected into fertilised one-cell stage embryos. Guide RNAs and targeting vector sequences are provided in Extended Data Table 2. Heterozygous knock-in mice were first backcrossed to C57BL/6J mice for 2 generations to eliminate potential off-target events. Gene-swap mice were maintained on a C57BL/6J background unless stated otherwise in the text (see below).

*Mcl-1^Bcl-xL^* and *Mcl-1^Bcl-2^* gene-swap mice on a C57BL/6J background were bred to F1 hybrids of the FVBxBALB/c strains, this F1 hybrid was chosen because these mice are among the most genetically diverse laboratory mouse strains, which improves fertility and vigor^54^.

The *RosaCreERT2^+/Ki^* and *Mcl-1^fl/fl^* mice, both on a C57BL/6J background, have been described previously^55,56^. To activate CreERT2, mice were given 90 mg/kg body weight tamoxifen (Sigma-Aldrich, #10540-29-1) in peanut oil/10% ethanol each day for 2 consecutive days by oral gavage.

Mice were weaned between 19-23 days of age and deemed adult at 8 weeks of age. Male and female animals were both used as they became available or occurred during dissection of embryos and foetuses. No animals and no data were excluded. All animal experiments were performed with the approval of the Walter and Eliza Hall Institute Animal Ethics Committee and according to the Australian code of practice for the care and use of animals for scientific purposes.

For the genotyping of adult mice and embryos, ear clip samples or embryo tail clip samples, respectively, were lysed in Direct PCR (Tail) lysis buffer (Viagen, Biotech, #102-T) supplemented with 40 µg/mL Proteinase K (Sigma-Aldrich, #EO0491). Genotyping was performed in-house using specific oligonucleotides listed in Extended Data Table 2.

### Pre-implantation embryo handling and analysis

For timed matings, noon of the day on which the vaginal plug was first observed was defined as embryonic day 0.5 (E0.5). Pre-implantation embryos were recovered from the oviduct at E0.5, E2.5 and from the uterus at E3.5.

E0.5 embryos were cultured in embryo culture medium (EmbryoMax® Human Tubal Fluid, Sigma-Aldrich, # MR-070-D) under mineral oil for embryo culture (Sigma-Aldrich, # M5310) to observe pre-implantation development *ex vivo* and photographed daily (Nikon Diaphot 300) equipped with a digital camera (Zeiss).

The zonae pellucidae were removed from E2.5 and E3.5 embryos using Acid Tyrode’s solution (0.8 g NaCl, 0.02 g KCl, 0.024 g CaCl_2_•2H_2_O, 0.01 g MgCl_2_•6H_2_O, 0.1 g glucose and 0.4 g polyvinylpyrrolidone dissolved in 100 mL of H_2_O).

For direct genotyping, zona-less E2.5 and E3.5 embryos were placed into 10 µL of DNA lysis buffer (1X MyTaq Red Mix, Bioline, #25044) complemented with 0.2 µL of 10 mg/mL proteinase K (Sigma-Aldrich, # EO0491) and incubated at 55°C for 2 h. The proteinase K was inactivated at 85°C for 1 h, before conducting the genotyping PCR.

Inner cell mass (ICM) outgrowth cultures were performed as reported previously^57^. In brief, E3.5 embryos were plated individually into gelatine-coated wells of 24-well plates in embryonic stem cell (ESC) medium (DMEM, high glucose 4500 mg/L (Gibco, ThermoFisher, #11965,), 100 µM 2-mercapto-ethanol (Sigma-Aldrich, #M-7522), 1 x non-essential amino acids (Gibco, ThermoFisher, #11140050), 1 x L-glutamine (Gibco, ThermoFisher, # 25030-024), 1 x sodium pyruvate (Gibco, ThermoFisher, #11360070,), 1000 IU LIF/mL (Sigma-Aldrich, #ESG1107) and 20% ESC-qualified fetal bovine serum (FBS; Gibco, ThermoFisher, #26140079)). Embryos were incubated at 37°C in 5% (v/v) CO_2_ in air and photographed daily using an inverted microscope (Nikon Diaphot 300) equipped with a digital camera (Zeiss). Cultures were lysed and genotyped after 7 to 9 days of culture.

Terminal deoxynucleotidyl transferase dUTP nick end labelling (TUNEL) staining of pre-implantation embryos was performed as reported previously^58^. In brief, zona-less E2.5 embryos were immobilised on gelatine-coated microscope slides by fixation in 1% (w/v) paraformaldehyde (Sigma-Aldrich, #P6148) in PBS, washed, post-fixed in cooled ethanol:acetic acid (2:1) and stained using a TUNEL kit (ApopTag® Fluorescein In Situ Apoptosis Detection Kit, Merck Millipore, #S7110) according to the manufacturer’s instructions. After photography using an upright microscope (Zeiss) and a digital camera (Zeiss), coverslips were removed by submerging slides in PBS, the embryos were individually scraped off the microscope slide using drawn-out glass pipettes (new pipette for each embryo) filled with 1X MyTaq Red mix (Bioline, #25044), aspirated and transferred into 10 µL of DNA lysis buffer and processed for genotyping as described above.

### Post-implantation embryo handling and analysis

For timed matings, noon of the day on which the vaginal plug was first observed was defined as embryonic day 0.5 (E0.5). Mouse embryos and foetuses were recovered at E10.5, E11.5, E12.5, E13.5, E14.5, E15.5 and E18.5 and examined and scored before genotyping (i.e., blinded to genotype). Embryos were imaged using the SV11 stereomicroscope from Zeiss equipped with the ZEN software (Zeiss). Embryo images were further processed using the Adobe Photoshop software. For some images the healing brush spot tool was used to delete image parts of neighbouring embryos from the background and scratches in the dish, but no such changes have been made to the embryos. Other changes, including level adjustment (black/white balance), brightness adjustment and exposure adjustment have been applied uniformly to the entire image. Embryos and data were only excluded when technical errors occurred (e.g., no clear genotyping result or damage during dissection).

### Cell lines and *in vitro* cell survival assays

Mouse embryonic fibroblasts (MEFs) were prepared from E11.5 embryos of the indicated genotypes. Briefly, the foetal liver and head were removed, and the remaining tissue was incubated for 5-10 min in trypsin to form a single cell suspension. Cells were washed with Dulbecco’s Modified Eagle Medium (DMEM, Gibco, ThermoFischer, #<u>12491015</u>) and seeded onto 0.1% gelatine-coated 6-well plates in DMEM supplemented with 10% foetal calf serum (Gibco, ThermoFischer, #26140079) and 1x Glutamax (Gibco, ThermoFischer, #35050061). Primary MEFs were cultured in a low oxygen (3%) incubator to prolong their lifespan. Where indicated, MEFs were immortalised with the SV40 Large T antigen. Cells were cultured in a 6-well plate and transfected with 10 µg of linearised plasmid expressing the SV40 Large T antigen using the FuGene Transfection Kit (Promega, #E2691) according to the manufacturer’s instructions. Cells were then serially passaged, with complete immortalisation observed at passage 8-10. Immortalised MEFs were cultured in DMEM supplemented with 10% foetal calf serum (Gibco, ThermoFischer, #26140079), 1x Glutamax (Gibco, Thermo Fischer #35050061), 50 µM 2-mercaptoethanol (Sigma-Aldrich, #M6250), and 100 mM asparagine (Sigma-Aldrich, #70-47-3). Cell survival assays were performed in 96-well flat bottom plates containing 5x10^4^ MEFs per well. Cells were treated with the ABT-199 (Active Biochem. #A-1231), A-1331852 (in house)^59^, S-63845 (Active Biochem, #6044), Etoposide (Sigma-Aldrich, #33419-42-0), Thapsigargin (Sigma-Aldrich, #T9033), Nutlin-3a (Selleckchem, #S8059) or Ionomycin (Sigma-Aldrich, #56092-82-1) at the concentrations indicated in the figures and figure legends for the indicated time periods. Cell viability was assessed by MTT assay (Roche, #11465007001) according to the manufacturer’s instructions.

Mitogen-activated B cells were generated from splenocytes isolated from adult mice. In brief, single cell suspension were generated by mashing the spleen through a 100 µm cell strainer. Cells were washed twice with PBS and 5x10^3^ cells/mL were plated in RPMI 1640 medium (Gibco, ThermoFischer, #11965) supplemented with 1x Glutamax (Gibco, ThermoFischer, #35050061), 50 µM 2-mercaptoethanol (Sigma-Aldrich, #M6250), 100 mM asparagine (Sigma-Aldrich, #70-47-3), non-essential amino acids (Gibco, Thermo Fischer, #11140050). B cells were activated by adding 100 units/mL IL-4 (in house), 10 µg/mL rat anti-CD40 (clone FGK45, in house), 10 ng/mL LPS (Lipopolysaccharides from *Escherichia coli*, Sigma-Aldrich, #L2630) to the culture for 48 h. Cell survival assays were performed in 96-well flay bottom plates containing 5x10^4^ cells per well. Cells were treated with ABT-199 (Active Biochem. #A-1231), A-1331852 (in house)^59^, S-63845 (Active Biochem, #6044) or Cytarabine (Pfizer, #C587PB) at the concentrations indicated in the figures and figure legends and cell survival was assessed at the indicated time points post-treatment by flow cytometry (BD Fortessa X20) after staining with annexin V conjugated to Alexa Fluor plus 647 (ThermoFischer, #P1304MP) and propidium iodide (PI, ThermoFischer, #P1304MP)). Viable cells were identified as those negative for both annexin V and PI.

### Western blot analysis

Total cell lysates were generated from E10.5 embryos using CHAPS buffer (10 mM HEPES, pH 7.4, 150 mM NaCl, 1% CHAPS plus complete protease inhibitor cocktail (Roche, #11697498001)). After determining the protein concentration of each sample using the Protein Assay Dye Reagent Concentrate (Biorad, #50000002)), 30 µg of lysate was resolved on 10%-14% NuPage Bis-Tris-polyacrylamide gels (Invitrogen) and proteins were then transferred onto PVDF/nitrocellulose membranes (Amersham). Blots were then incubated with blocking buffer (Tris Buffered Saline, 0.1% Tween20, 5% milk) for 1 h at room temperature and probed with the primary antibodies at 4°C ON. A list of the antibodies used is provided in Extended Data Table 3. Incubation with HRP-conjugated secondary antibodies was subsequently performed for at least 1 h, and protein bands were visualised on the ChemiDoc Touch Imaging System (Biorad) using the Luminata Forte enhanced chemiluminescence reagent (Millipore, #RPN2209)). Western blot images were initially analysed using the ImageLab software (Biorad) and further processed using the Adobe Photoshop software, including image cropping and brightness adjustment that were applied uniformly to the entire image.

Positive controls for specific antibodies were used. These included the murine AML cell line MLL-AF9 (expressing high levels of mBCL-XL), the human DLBCL cell line DOHH2 (expressing high levels of hBCL-2), the murine *Eµ-Myc* double-hit lymphoma (DHL) cell lines 214DHL and 216DHL (expressing high levels of mBCL-2 and mMCL-1, respectively) and the murine *Eµ-Myc* lymphoma cell line AH15A (expressing high levels of mBAX and mBAK).

### Mitochondrial isolation and blue native-page analysis

Crude mitochondria were isolated from cultured cells as previously described^60^. MEFs were seeded onto 150 mm tissue culture dishes (Corning, #CLS430599) and cultured to confluency in DMEM (Gibco, #11885084; Thermo Fisher Scientific) supplemented with 10% foetal calf serum, 1 x penicillin/streptomycin (Sigma-Aldrich; #P4458),1x GlutaMAX (Life Technologies; #35050061) and 100 mM L-asparagine (Sigma-Aldrich; #A4159). Cells were dissociated from culture dishes using PBS + 2 mM EDTA and centrifuged at 800g for 5 min at 4°C. Supernatants were aspirated and cell pellets washed in cold PBS and centrifuged at 800g for 5 min at 4°C. Supernatants were aspirated and cell pellets frozen at -80°C for at least 30 min. To isolate the crude mitochondrial fraction, thawed cell pellets MEFs and primary hepatocytes were resuspended in Isolation Buffer A (20 mM HEPES-KOH pH 7.6, 220 mM mannitol, 70 mM sucrose, 1 mM EDTA, 0.5 mM PMSF and 2 mg/mL BSA) and lysed using 20 strokes of a Dounce glass homogenizer. Cell lysates were centrifuged at 800g for 5 min at 4°C to pellet nuclei and remaining intact cells.

Supernatants containing mitochondria were transferred into clean tubes and centrifuged at 13,000g for 10 min at 4°C. Supernatants were aspirated and the remaining mitochondrial pellets were resuspended in Isolation Buffer B (20 mM HEPES-KOH pH 7.6, 220 mM mannitol, 70 mM sucrose, 1 mM EDTA and 0.5 mM PMSF) and centrifuged again at 13,000*g* for 10 min at 4°C. Supernatants were aspirated and mitochondrial pellets were resuspended in 500 μL Isolation Buffer B if used immediately or 500 μL Sucrose Buffer (10 mM HEPES pH 7.6 and 0.5 M sucrose) if aliquoted and frozen for later use. Protein concentration was determined by using the bicinchoninic assay (BCA; ThermoFisher, #23223). BN-PAGE was performed as previously described ^61^ with continuous 4%-13% gels made using a gradient mixer, the separating gels composed of acrylamide/bis-acrylamide solutions made in BN gel buffer (66 mM ε-amino n-caproic acid, 50 mM Bis-Tris pH 7.0) with 13% acrylamide/bis-acrylamide and 20% (w/v) glycerol or 4% acrylamide/bis-acrylamide alone) and the stacker gel composed of 4% acrylamide/bis-acrylamide in BN gel buffer and layered onto the separating gel. Crude mitochondria were pelleted from the solution at 13,000g, 10 min at 4°C, with supernatants subsequently removed and the remaining mitochondrial pellets solubilised in digitonin detergent buffer (20 mM Bis-Tris pH 7.0, 50 mM NaCl, 10% (w/v) glycerol and 1% digitonin) for 10 min on ice, followed by centrifugation at 13,000g for 10 min at 4°C. The supernatants containing solubilised mitochondrial complexes were then transferred into a clean tube containing one-tenth the volume of 10X BN-PAGE loading dye (5% (w/v) Coomassie blue G250, 500 mM ε-amino n-caproic acid, 100 mM Bis-Tris pH 7.0) prior to gel loading. Gel electrophoresis was performed using BN-PAGE blue cathode buffer (50 mM tricine, 15 mM Bis-Tris, 0.02% (w/v) Coomassie blue G250), which was replaced with BN-PAGE clear cathode buffer (50 mM tricine, 15 mM Bis-Tris) once the dye front has migrated halfway through the gel, and BN-PAGE anode buffer (50 mM Bis-Tris pH 7.0). Transfer to PVDF membranes (Merck, #IPVH00010) was performed using an Invitrogen Power Blotter System (Thermo Fisher) according to the manufacturer’s instructions. Primary antibodies, sources and associated dilutions are listed in the Extended Data Table 3. Anti-mouse IgG (Sigma-Aldrich, #A9044) or anti-rabbit IgG (Sigma-Aldrich; A0545) antibodies conjugated to horseradish peroxidase were used as secondary reagents at a dilution of 1:10,000. Clarity Western ECL chemiluminescent substrate (BioRad; #1705061) was used for detection on a BioRad ChemiDoc XRS+ imaging system according to the manufacturer’s instructions.

### Whole-mount embryo labelling and imaging

Mouse embryos were recovered at E12.5 and fixed in 4% (w/v) paraformaldehyde (Sigma-Aldrich, #P6148)/PBS at room temperature (RT) over night (ON). Antibody labelling and clearing were performed following the iDISCO protocol (https://www.cell.com/fulltext/S0092-8674(14)01297-5). In brief, embryos were washed 3x with PBS for 30 min and dehydrated with a methanol/H_2_O series (20%, 40%, 60%, 80%, 100% (2x)) for 1 h for each dilution at room temperature. Embryos were incubated overnight in 66% dichloromethane (DCM, Sigma-Aldrich, #75-09-2)/33% methanol at room temperature and washed 2x in 100% methanol at room temperature and then chilled at 4°C. Embryos were bleached in chilled fresh 5% H_2_O_2_ in methanol ON at 4°c and then rehydrated with a methanol/H_2_O series (80%, 60%, 40%, 20%, PBS) for 1 h each at room temperature. Embryos were washed twice with PTx (PBS/0.2% Triton-X-100) for 1 h at room temperature, incubated with permeabilization buffer (PTx/20%DMSO/38 mg/mL glycine) for 1 day and blocking buffer (PTx / 10%DMSO / 6% donkey serum) for 2 days. For immunolabelling embryos were washed 4 times with PTwH (PBS/0.2% Triton-X-100/10 µg/mL heparin (Stemcell Technologies, #07980)) for 1 h each at 37°C and incubated with primary antibody (PE-labelled rabbit anti-CD31/PCAM1 (1:100, cloneC31.7, Abcam, #ab215753) in PTwH/5% DMSO/3% Donkey Serum (Abcam, #7475) for 5 days at 37°. For clearing, embryos were washed 4 times with PTwH for 1 h each at 37°C and then dehydrated with a methanol/H_2_O series: 20%, 40%, 60%, 80%, 100% (2x); for 1 h each dilution at room temperature. On the next day, embryos were incubated in 66% DCM/33% methanol for 3 h at room temperature. Embryos were washed with 100% DCM 3x for 15 min at room temperature and incubated DiBenzyl Ether (DBE, Sigma-Aldrich, #103-50-4) for 15 min. Cleared and stained embryos were transferred into ethyl cinnamate (Sigma-Aldrich, #103-36-6). Light sheet imaging was performed using a Zeiss Z.1 Light sheet microscope equipped with a 5x/0.16 detection objective. 3D tiled image stacks of autofluorescence (excitation 405 nm, emission 505-545 nm) and PE fluorescence (excitation 561 nm; emission 575-615 nm) were acquired sequentially at a voxel size of XX in XY and XX in Z. Data sets were 3D reconstructed using Imaris v9.7.1 (Bitplane). Imaris version 9.7.1 (Bitplane) was used for image processing.

### Immunofluorescence analysis of adult liver sections

Liver samples were harvested, fixed and embedded in paraffin as described above. Histological sections (5 µm) on slides were de-waxed and heat-induced epitope retrieval was performed using Tris-EDTA (10 mM Tris, 1 mM EDTA) pH 9.0 retrieval buffer. Slides were washed 2-3 times with tap water followed by 2 washes with PBS containing 0.1% Triton X-100 (“0.1% PBS-T”). Next, sections were blocked for 1 h at room temperature using 10% horse serum (ThermoFisher, # 26050088) and 2% mouse-on-mouse (M.O.M) IgG blocking reagent (Vector Labs) diluted in 0.1% PBS-T. Sections were washed 3 times with PBS containing 0.01% Triton X-100 (“0.01% PBS-T”). This was followed by the application of primary antibodies (diluted in 0.01% PBS-T containing 1% horse serum), which were incubated overnight at 4°C. The following primary antibodies were used: mouse anti-Glutamine Synthetase (1:200,), rabbit anti-MCL-1 (1:100) (details for primary antibodies are listed in Extended Data Table 3). The next morning, sections were washed 3 times with 0.01% PBS-T before incubating for 1 h at room temperature with secondary antibodies (also diluted in 0.01% PBS-T containing 1% horse serum). The following secondary antibodies were used: donkey anti-mouse IgG conjugated to AlexaFluor plus 488 (1:500, ThermoFisher, #32766), donkey anti-rabbit IgG conjugated to AlexaFluor plus 647 (1:500, ThermoFisher, #A32795). After incubation with secondary antibodies, sections were washed 3 times with 0.01% PBS-T and DAPI (Sigma-Aldrich, #28718-90-3) nuclear stain, diluted in PBS, was applied for 15-20 min at room temperature. Subsequently, sections were washed 3 times with 0.01% PBS-T, mounted with Fluoromount-G (ThermoFisher, #00-4958-02) mounting medium and stored in the dark at 4°C until ready for imaging. Sections were imaged on a Zeiss LSM 980 confocal microscope and images processed using FIJI (v2.9.0).

### Electron microscopy

Cells were fixed in 2.5% glutaraldehyde in 0.1 M cacodylate buffer for 1 h at room temperature followed by 3x 10 min rinse in 0.1 M cacodylate buffer. Cells were osmicated in 1% osmium tetroxide in 0.1 M cacodylate buffer for 30 min at room temperature which was then reduced with 1.5% potassium ferrocyanide in 0.1 M cacodylate buffer for 30 min at room temperature. After rinsing with mQH_2_O (3x10min at room temperature) the cells were stained with 2.5% uranyl acetate (aqueous) overnight at 4°C. Cells were then rinsed as previously (mQH_2_O for 3x10mins at room temperature) and scrapped and pelleted for embedding in agarose (4% low melting point agarose, aqueous). Agarose blocks were de-hydrated with increasing concentrations of ethanol before final dehydration with 100% acetone (ethanol: 20%, 50%, 70%, 90%, 2 x 100%; acetone: 2 x 100%), aided by a microwave regime for 40 sec for each step (150 W, no vacuum, Pelco Biowave). Agarose blocks were then infiltrated with increasing concentrations of epon resin in acetone for 3 min for each step (25%, 50%, and 75%, 2 x 100%). Samples were left in fresh 100% resin overnight. After being transferred into silicone molds, blocks were polymerised in the oven at 60°C for 48 h. 70 nm sections were cut on a Leica UC7 ultramicrotome with a diamond knife (Diatome) and 150 mesh hexagonal copper grids. Transmission electron microscopy imaging was acquired on a Jeol JEM 1400-Plus at 80 kv. A combination of single snapshots and image montages were collected of cells using Jeol acquisition software (TEM Centre).

### Histology

Whole embryos and foetuses were sacrificed by cooling and then placed into 4% (w/v) paraformaldehyde (Sigma-Aldrich, #P6148)/PBS (E10.5-E14.5) or Bouin’s fixative (Australian Biostain, #ABF2.5l) (E18) and subsequently embedded in paraffin. E10.5-E13.5 Embryos were embedded in 4% low-melt point agarose (Sigma-Aldrich, #A9414) before paraffin embedding. 5 µm serial transverse sections were obtained and stained with haematoxylin and eosin (H&E). Stained slides were scanned using a 20x brightfield slide scanner (Polaris) and examined and imaged using the CaseViewer Software (3DHISTECH) or the QuPath software.

To examine tissues from pups and mice, samples of the brain, heart, kidney, liver, small intestine, large intestine, caecum and colon were dissected, fixed in 10% formalin and embedded in paraffin. Histological sections (5 µm) were subsequently stained with H&E or cresyl violet acetate (Sigma-Aldrich, #C5042). Immunohistochemistry to detect activated (cleaved) caspase-3 was performed using a specific antibody detecting cleaved caspase-3 (Asp175, Cell Signaling, #9661). TUNEL assays were performed using the TUNEL detection kit (Abcam, #ab206386) according to the manufacturer’s instructions. To detect fibrosis, formalin-fixed heart sections were stained with Masson’s trichrome (Sigma-Aldrich, #HT15). To detect lipid accumulation, frozen sections were stained with OilRedO (Sigma-Aldrich, #1320-06-5). Stained slides were scanned using a 20x brightfield slide scanner (Polaris) and examined and imaged using the CaseViewer Software (3DHISTECH) or the QuPath software.

### Quantification and statistical analysis

Genotype distributions were analysed by computing the cumulative binomial probability of being less or equal to the expected value (pbinom) using R (version 4.2.2, 2022-10-31, The R Foundation for Statistical Computing).

Kaplan Meyer survival curves were analysed by Log-Rank (Mantel Cox) test using statistics software (Prism 9 for Mac, version 9.4.1; GraphPad).

For *in vitro* cell survival analysis, comparison of the percentages of cell survival between different genotypes and treatments were analysed using two-way ANOVA with subsequent Tukey’s multiple comparison analysis (comparing to the wild-type samples) using statistics software (Prism 9 for Mac, version 9.4.1; GraphPad). Body weight data were compared by two-tailed, unpaired T-test using statistics software (Prism 9 for Mac, version 9.4.1; GraphPad). For each experimental approach, the number of replicates (n) is defined as number of animals examined or repeat experiments performed and this is stated in the figures and/or figure legends. Data are presented as mean ±SEM as indicated in the figure legends.

## Acknowledgements

The authors thank WEHI Bioservices, particularly Giovanni Siciliano, Jaclyn Gilbert, Tracey Baldwin, Louise Spencer, Jessica Martin, Tom Kitson, Katie Franks, Kelly Trueman and Lauren Whelan for animal husbandry and help with experiments using mice; Bruno Helbert, Rainbow Chan and Natalie Coleman for genotyping; the WEHI Histology Centre, particularly Ellen Tsui, Emma Pan and Andrew Spence, and Lachlan Whitehead for imaging data processing. This work was supported by the National Health and Medical Research Council (NHMRC): program grant #101671 to AS and PB; fellowships #1020363 to AS, #1176789 to AKV, #2017971 to MJH; ideas grant #2021510 to KB and KMcA; project grant #1160618 to TT; the NMRC OF-IRG #MOH-000614 to NF and a fellowship from the Bodhi Education Fund to GD. The generation of the gene-swap mice used in this study was supported by Phenomics Australia and the Australian Government through the National Collaborative Research Infrastructure Strategy (NCRIS) program.

## Author Contributions

KB, MJH and AS conceived this study. KB, KMcA, MJH, TT, AKV and AS planned experiments and interpreted data. KB, KMcA, AK, AT, LG, CR, SM, VM, PA, TT and AKV performed experiments. GD, PB, TP and NF made critical interpretations and provided ideas. KB, AKV and AS wrote the manuscript and all co-authors edited it.

## Competing Interest Declaration

KB, AK, AT, SM, KM, VW, LG, PA, NF, TP, GD, PB, AKV, TT, MJH and AS are or were for some time employees of The Walter and Eliza Hall Institute. The Walter and Eliza Hall Institute receives and milestone payments and royalties from the sale of Venetoclax, parts of which are distributed to current and former employees. MJH and AS collaborated with Servier on the development of MCL-1 inhibitors and received financial support for some of their research.

## Address for correspondence

Kerstin Brinkmann and Andreas Strasser

The Walter and Eliza Hall Institute of Medical Research

1G Royal Parade, Parkville, Victoria 3052, Australia

Phone: +61-3-9345-2624, FAX: +61-3-9347-0852

---

**Extended Data Fig. 1.**
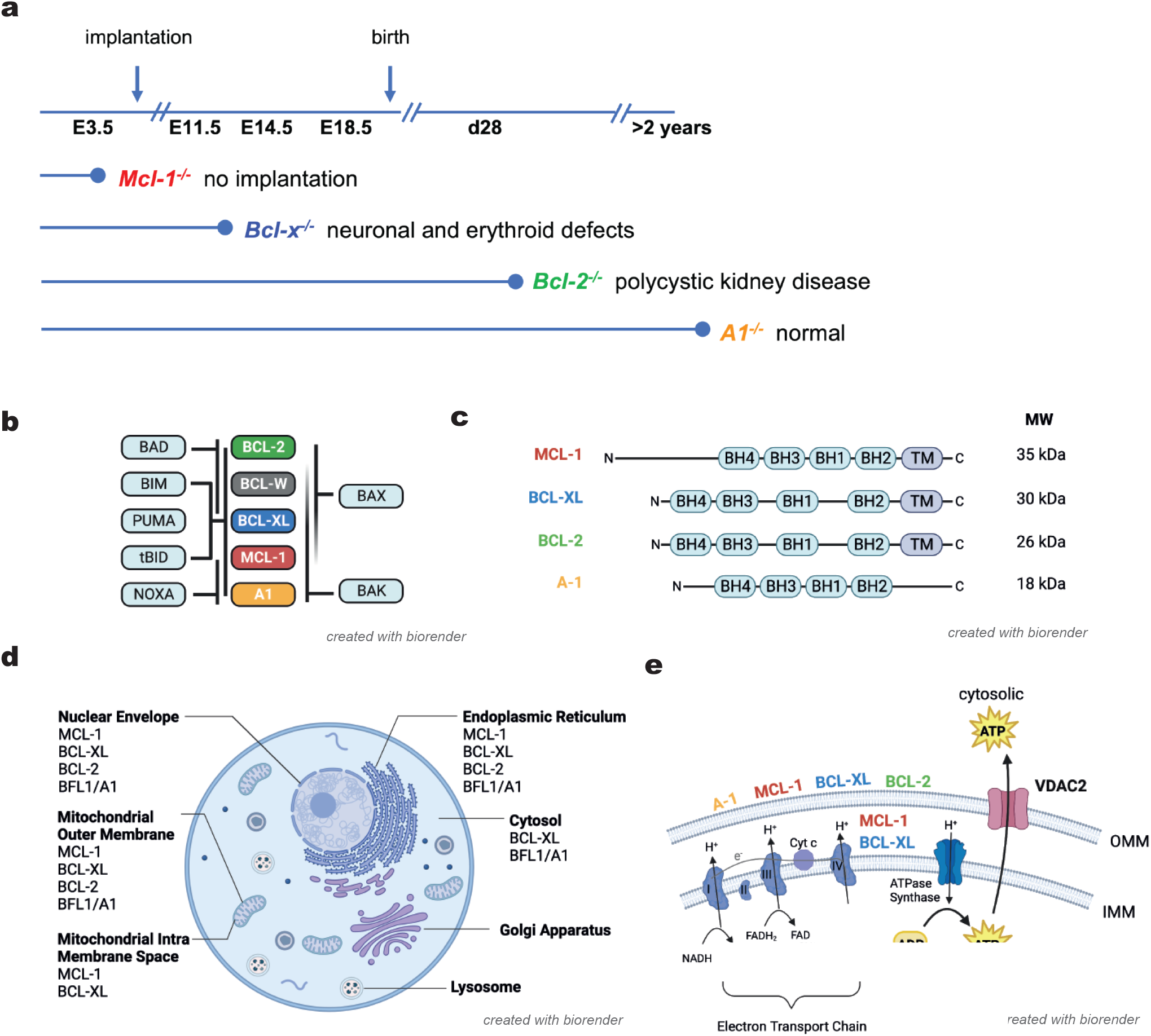
Similarities and differences of the anti-apoptotic members of the BCL-2 protein family. **a,** Schematic presentation of the time of developmental lethality and survival of *Mcl-1^-/-^*, *Bcl-x^-/-^*, *Bcl-2^-/-^* and *A1^-/-^* mice. **b**, Schematic presentation of the reported interactions of pro-apoptotic and anti-apoptotic members of the BCL-2 protein family. **c**, Schematic presentation of the protein domains of the anti-apoptotic BCL-2 family members, BCL-2, BCL-XL, MCL-1 and A1. The respective molecular weights (MW) of these proteins are indicated. **c**, Schematic presentation of the subcellular localisation of the anti-apoptotic BCL-2 proteins, BCL-2, BCL-XL, MCL-1 and A1. **d**, Schematic presentation of the localisation of the anti-apoptotic BCL-2 family members, BCL-2, BCL-XL, MCL-1 and A1 at the outer mitochondrial membrane (OMM) and in the space between the two mitochondrial membranes. e, Schematic presentation of the developmental stage and age of lethality of *Mcl-1^-/-^*, *Bcl-x^-/-^*, *Bcl-2^-/-^* and *A1^-/-^* mice.

**Extended Data Fig. 2.**
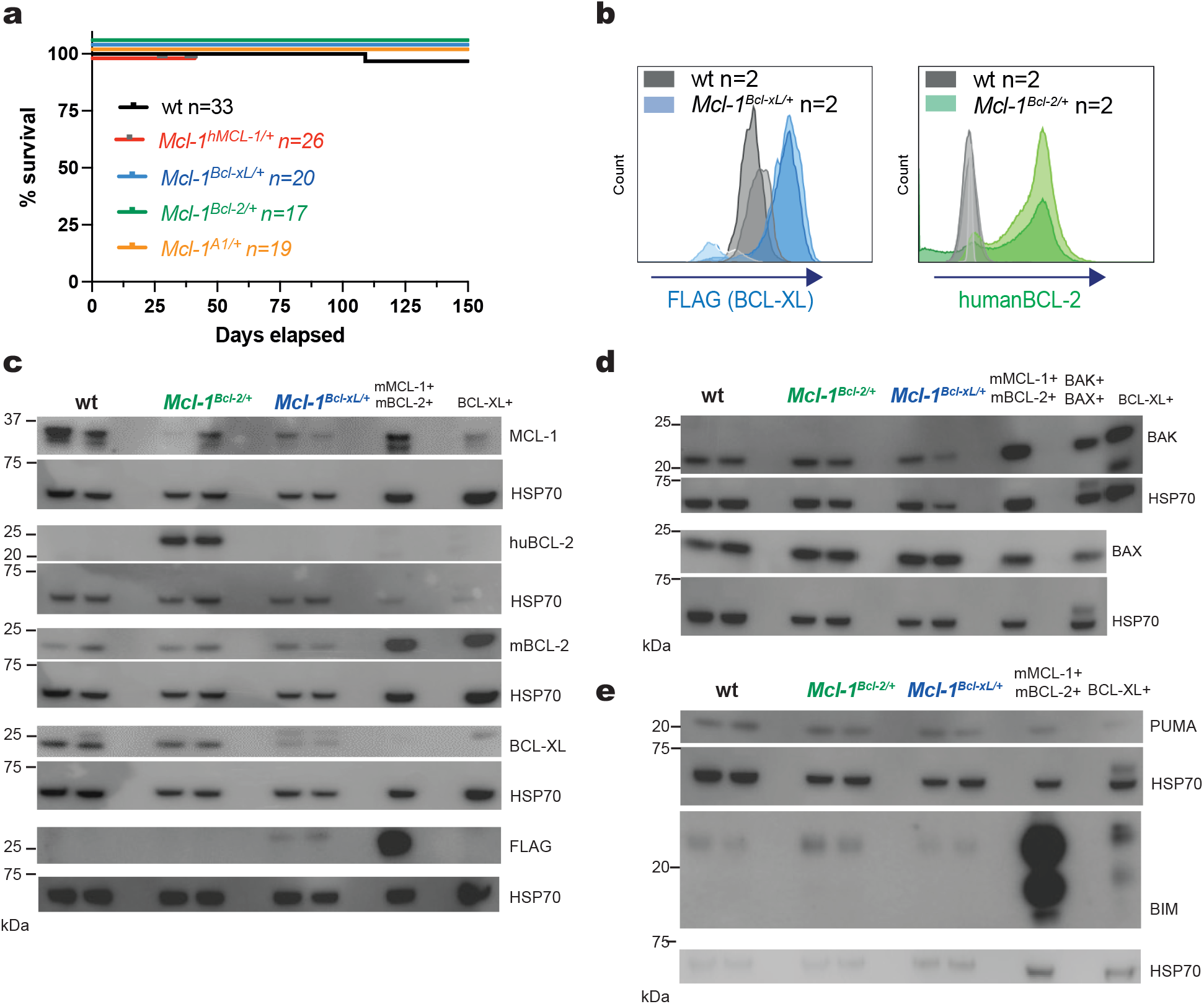
Expression analysis of the knock-in proteins in cells from heterozygous gene-swap mice. **a**, Kaplan-Meyer survival curves showing the survival of wild-type (n=33) and heterozygous *Mcl-1^hMcl-^*^1^*^/+^* (n=26), *Mcl-1^Bcl-xL/+^* (n=20), Mcl-1^Bcl-2/+^ (n=17) and *Mcl-1^A1/+^* (n=19) mice. Statistical significance was tested using the Log Rank (Mantle-Cox) test. No significant differences were observed. **b**, Flow cytometric analysis of the expression of FLAG-BCL-XL and human BCL-2 in bone marrow cells from wild-type and heterozygous *Mcl-1^Bcl-xL/+^* and *Mcl-1^Bcl-2/+^* gene-swap mice using monoclonal antibodies that specifically detect FLAG or human BCL-2, respectively (n=2 mice per genotype). **c-e**, Western blots of total protein lysates of primary mouse embryo fibroblasts (MEFs) that had been generated from E11.5 wild-type, *Mcl-1^Bcl-xL/+^* and *Mcl-1^Bcl-2/+^* embryos (n=2 per genotype, MEF isolates of individual embryos were loaded into individual lanes) detecting **c**, mouse MCL-1, mouse BCL-XL, FLAG-BCL-XL, human and mouse BCL-2, **d**, BAX and BAK and **e**, PUMA and BIM. Probing for HSP70 served as a protein loading control. The following positive controls were used: the murine AML cell line MLL-AF9 (expressing high levels of mBCL-XL), the human DLBCL cell line DOHH2 (expressing high levels of hBCL-2), the murine Eµ-Myc double-hit lymphoma (DHL) cell lines 214DHL and 216DHL (expressing high levels of mBCL-2 and mMCL-1, respectively), the murine Eµ-Myc lymphoma cell line AH15A (expressing high levels of BAX and BAK).

**Extended Data Fig. 3.**
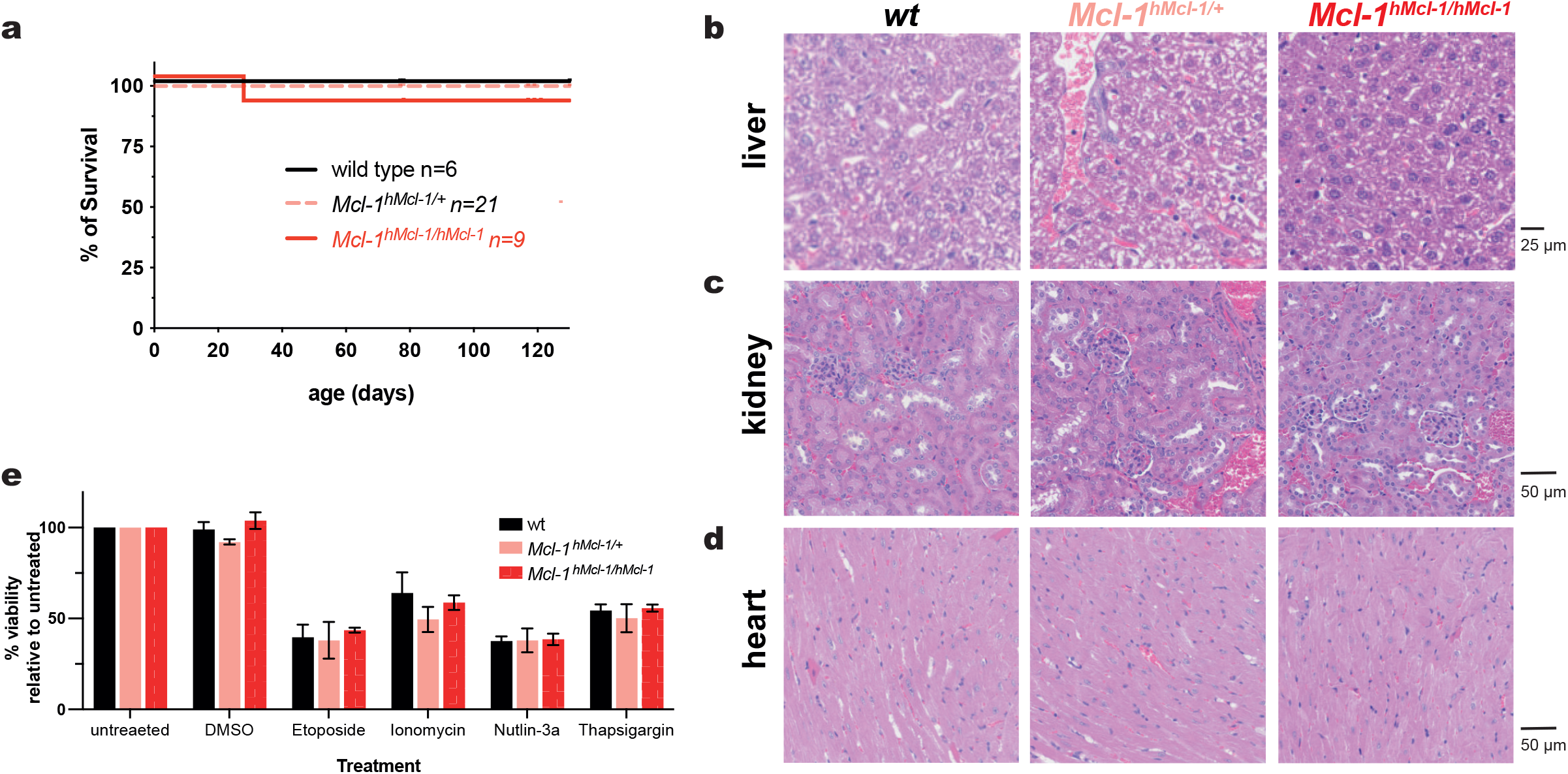
*Mcl-1^hMcl-1/hMcl-1^* control gene-swap mice are indistinguishable from wild-type mice. **a**, Kaplan-Meyer survival curves showing the survival of wild-type (n=6), *Mcl-1^hMcl-1/+^* (n=21) and *Mcl-1^hMcl-1/hMcl-1^* (n=9) mice. Statistical significance was tested using the Log Rank (Mantle-Cox) test. No significant differences were observed. **b-d**, Representative H&E-stained sections of the **b**, liver **c**, kidney and **d**, heart from wild-type (left panel), heterozygous *Mcl-1^hMcl-1/+^* (middle panel) and homozygous *Mcl-1^hMcl-1/hMcl-1^* (right panel) mice (50-60 days old). **e**, Survival analysis of primary mouse embryo fibroblasts (MEFs) generated from wild-type, heterozygous *Mcl-1^hMcl-1/+^* and homozygous *Mcl-1^hMcl-1/hMcl-1^* E11.5 embryos (n=2 per genotype). Cell viability was assessed 48 h after treatment with DMSO (ctrl), etoposide (20 µg/mL), ionomycin (2.5 µg/mL), the MDM2 inhibitor nutlin-3A (20 µM), thapsigargin (500 pM) by MTT assay. Data are presented as mean ± SEM from 2 independent experiments, each performed in triplicates. Statistical significance was tested using two-way ANOVA with Tukeys’s multiple comparison. No significant differences were observed.

**Extended Data Fig. 4.**
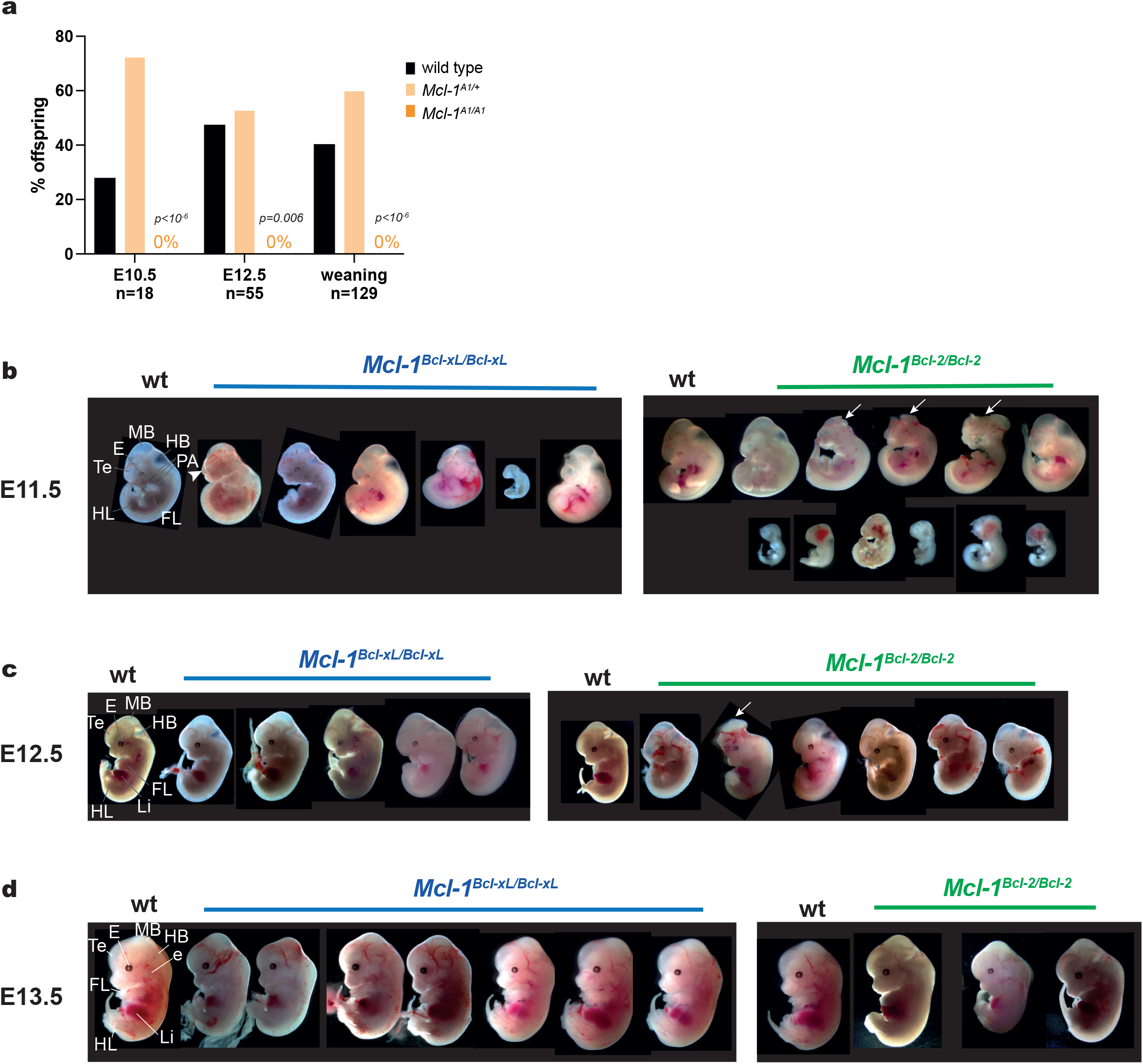
Assessment of *Mcl-1^Bcl-xL/Bcl-xL^* and *Mcl-1^Bcl-2/Bcl-2^* gene-swap embryos. **a**, Genotype frequency at E10.5 (n=18), E12.5 (n=25) and weaning (n=129) among offspring of intercrosses of *Mcl-1^A1/+^* mice. Datasets for the offspring distribution at E10.5 are also presented in Fig. 1f and are included here again for comparison. Genotype distributions were analysed by computing the cumulative binomial probability of being less or equal to the expected value. p-values are indicated in the figure. **b-d,** Representative images showing wild-type, homozygous *Mcl-1^Bcl-xL/Bcl-xL^* and homozygous *Mcl-1^Bcl-2/Bcl-2^* embryos at embryonic day **b**, E11.5, **c**, E12.5, **d**, E13.5. **Labelling**: E, eye; e, ear; FL, forelimb (forelimb bud, E11.5 and E12.5); HB, hindbrain; HL, hindlimb (hindlimb bud, E11.5 and E12.5); Li, liver; MB, midbrain; PA, pharyngeal arches; Te, telencephalon. Arrowhead indicates collapsed telencephalic vesicle. Arrows indicate exencephaly.

**Extended Data Fig. 5.**
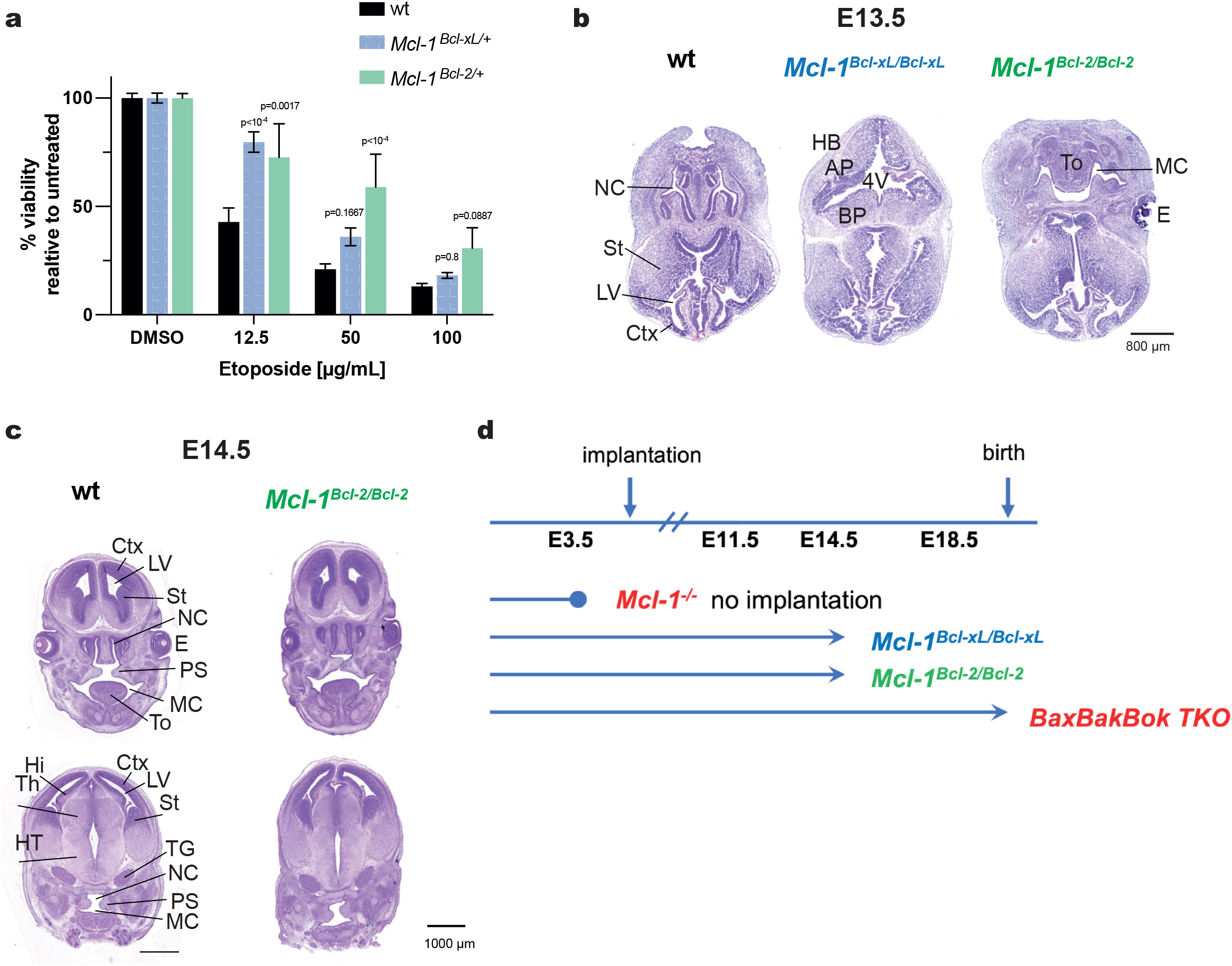
Decreased apoptosis in *Mcl-1^Bcl-xL^* and *Mcl-1^Bcl-2^* gene-swap embryos. **a**, Survival of primary mouse embryo fibroblasts (MEFs) generated from wild-type, *Mcl-1^Bcl-xL/+^* and *Mcl-1^Bcl-2/+^* E11.5 embryos (n=2 per genotype). MEFs were treated for 48 h with the indicated concentrations of etoposide. Cell viability was assessed by MTT assay. Data are presented as mean ±SEM from 2 independent experiments, each performed in triplicates. Statistical significance was tested using two-way ANOVA with Tukey’s multiple comparison. p-values are indicated in the figure. **b**, Representative images of H&E-stained serial sections of the head region of homozygous *Mcl-1^Bcl-xL/Bcl-x^*^L^ and homozygous *Mcl-1^Bcl-2/Bcl-2^* embryos at embryonic day E13.5. **c**, Representative images of H&E-stained serial sections of the head region of homozygous *Mcl-1^Bcl-2/Bcl-2^* embryos at embryonic day E14.5. Images of wild-type littermates are shown for comparison. **d**, Schematic presentation of the time of developmental lethality and survival of *Mcl-1^-/-^*, *Mcl-1^Bcl-xL/Bcl-xL^*, *Mcl-1^Bcl-2/Bcl-2^* and *Bax^-/-^Bak^-/-^Bok^-/-^* mice. **Labelling**: 3V, third ventricle; 4V, mesencephalic vesicle (future 4th ventricle); AP and BP, alar and basal plate of the hindbrain; Ctx, developing cortes; E, eye; HB, hindbrain; Hi, developing hippocampus; HT, hypothalamus (diencephalon); LV, telencephalic vesicle or lateral ventricle; MC, mouth cavity; NC, nasal cavity; PS, palatal shelf, here elevated; St, striatum; TG, trigeminal ganglion; Th, thalamus (diencephalon); Te, telencephalon; To, tongue. Arrows indicate collapsed telencephalic and mesencephalic vesicles. Arrowheads indicate neuroepithelial folds that would appear as ridges when external morphology was examined.

**Extended Data Fig. 6.**
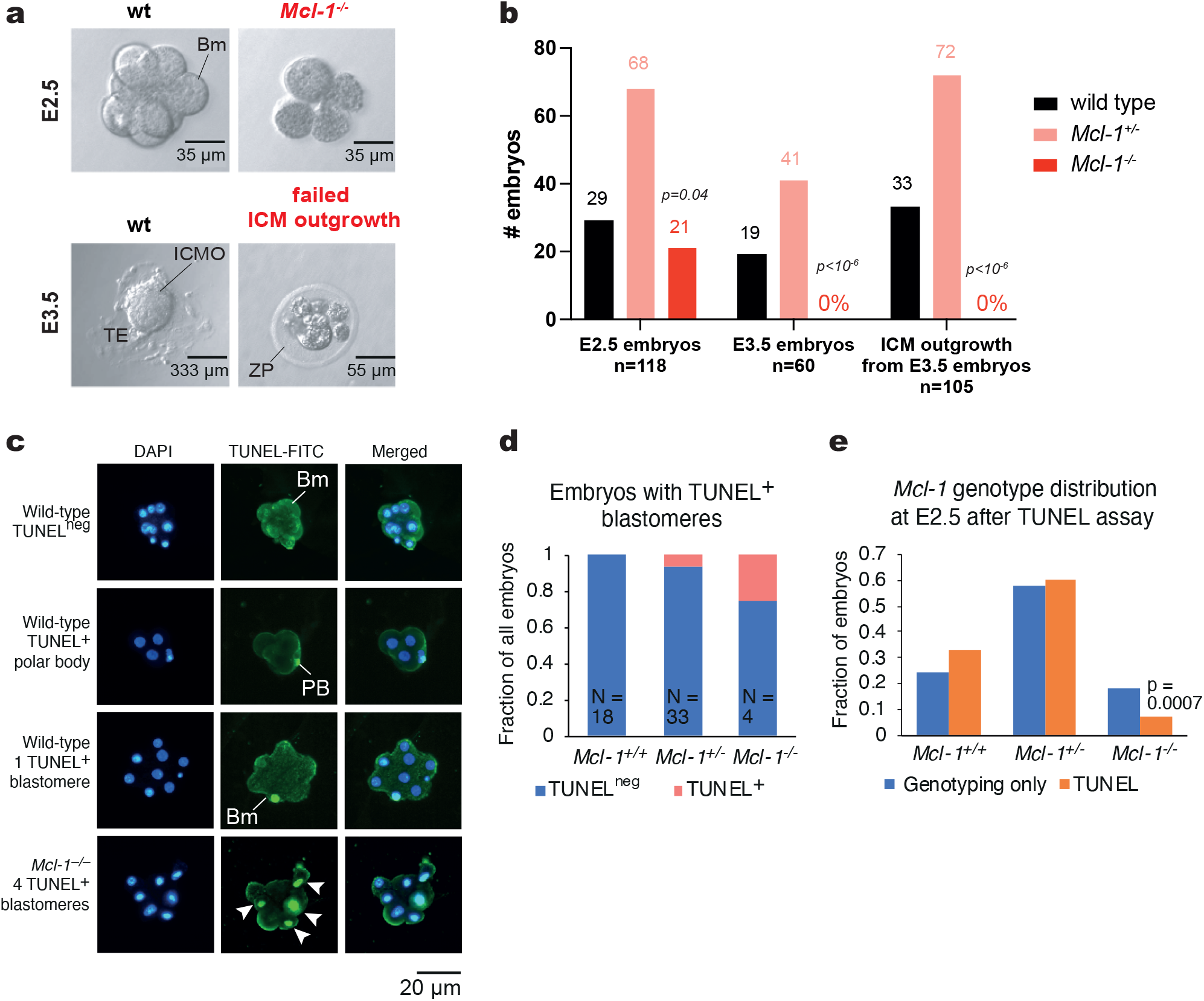
*Mcl-1^-/-^*E2.5 embryos (FVBxBALB/cxC57BL/6) die due to increased apoptosis. **a**, Representative images showing wild-type and *Mcl-1^-/-^* E2.5 embryos (FVBxBALB/cxC57BL/6 mixed genetic background, upper panels). The inner cell mass (ICM) outgrowth was imaged after 5 days of culture of E3.5 blastocysts (lower panel). **b**, Genotype frequencies at E2.5 and E3.5 and ICM outgrowths from E3.5 embryos of intercrosses of *Mcl-1^+/-^* mice (FVBxBALB/cx C57BL/6 mixed genetic background). Genotype distributions were analysed computing the cumulative binomial probability of being less or equal to the expected value. p-values and number of embryos (n) are indicated in the figure. **c**, Immunofluorescence images of TUNEL staining analysis in *Mcl-1^+/+^*and *Mcl-1^-/-^* E2.5 embryos (FVBxBALB/cxC57BL/6 mixed genetic background) counterstained with DAPI (blue). Embryos were recovered from slides and subjected to genotyping by genomic PCR after TUNEL staining and photography. 1st row, negative, background signal (dull green, non-nuclear); 2nd row, TUNEL positive polar body (bright green, small, no cytoplasm); 3rd row, embryo with a single TUNEL positive blastomere (bright green, large with cytoplasm compared to polar body); 4th row, *Mcl-1^-/-^*embryo with four TUNEL positive blastomeres. **d**, Genotype frequency among E2.5 embryos recovered from intercrosses of *Mcl-1^+/-^* mice (FVBxBALB/cxC57BL/6 mixed genetic background), either directly after recovery (genotyping only) or after TUNEL staining (TUNEL). Note the loss of *Mcl-1^-/-^*embryos compared to the direct genotyping (p=0.0007), suggesting that *Mcl-1^-/-^* embryos failed to adhere to the slides during the TUNEL procedure (detecting DNA fragmentation). Genotype distributions were analysed computing the cumulative binomial probability of being less or equal to the expected value. p-values are indicated in the figure. **e**, The fraction of embryos with TUNEL positive blastomeres among *Mcl-1^+/+^* (n=18), *Mcl-1^+/-^* (n=33) and *Mcl-1^-/-^*(n=4) embryos (FVBxBALB/cxC57BL/6 mixed genetic background). **Labelling**: Bm, blastomere; ICMO, inner cell mass outgrowth; TE, trophectoderm; ZP, zona pellucida; PB, polar body. Arrowheads indicate TUNEL positive blastomeres in the *Mcl-1^-/-^* embryo.

**Extended Data Fig. 7.**
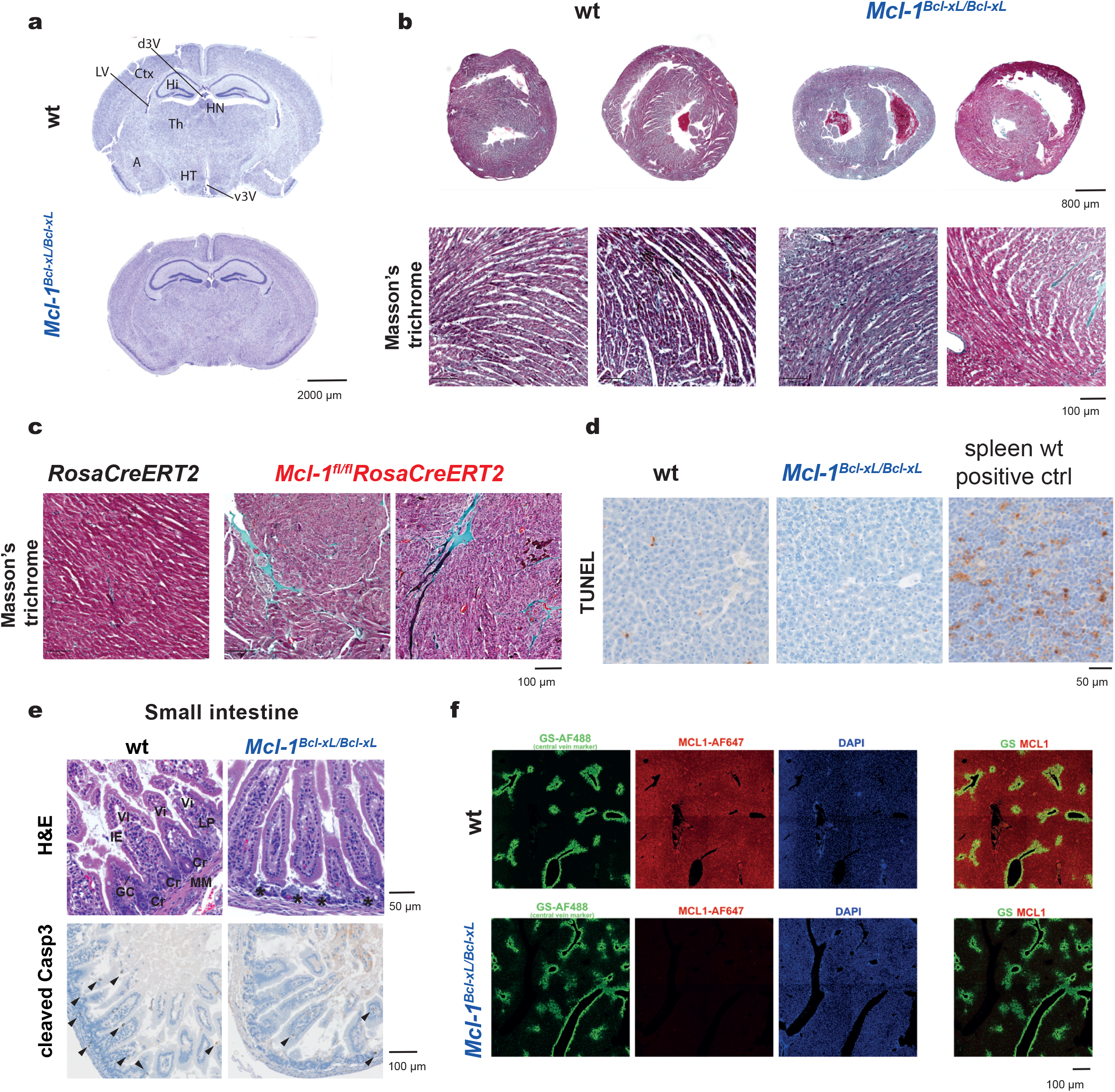
Loss of the non-apoptotic function of MCL-1 is critical for survival of mice after birth. **a**, Representative images of cresyl violet-stained coronal serial sections of the brains from 21-day-old wild-type (wt) (top) and homozygous *Mcl-1^Bcl-xL/Bcl-xL^*(bottom) mice (FVBxBALB/ cxC57BL/6 genetic background). **b**, Representative images of Mason’s trichrome-stained sections (detecting lipids) of the heart (myocardium) from 21-day-old wild-type (wt n=2, left panels) and homozygous *Mcl-1^Bcl-xL/Bcl-xL^* (n=2, right panels) mice (FVBxBALB/cxC57BL/6 genetic background). **c**, Representative images of Masson’s trichrome-stained sections (detecting lipids) of the heart (myocardium) from 10-12 weeks old-old *RosaCreERT2* (n=1, left panel) and *Mcl-1^fl/fl^RosaCreERT2* mice (n=2, right panels) 48 h after tamoxifen induced CRE activation and deletion of floxed alleles. **d**, Representative images of TUNEL stained sections from livers of 21-day-old wild-type and homozygous *Mcl-1^Bcl-xL/Bcl-xL^* mice (FVBxBALB/cxC57BL/6 genetic background). Spleen sections form wild-type mice are shown as a positive control; a few TUNEL positive cells are readily detectable in the control tissue. **e**, Representative histological images of H&E-stained sections (top row) and sections stained for cleaved (activated) caspase3 (bottom row) of wild-type (left panel) and homozygous *Mcl-1^Bcl-xL/Bcl-xL^* mice (right panel) (FVBxBALB/cxC57BL/6 mixed genetic background) of the small intestine. **f**, Representative immunofluorescence imaging of liver sections from 21-day-old wild-type (wt) and homozygous *Mcl-1^Bcl-xL/Bcl-xL^*mice (mixed FVBxBALB/cxC57BL/6 genetic background) showing the expression of the central vein marker glutamine synthetase (GS, green), MCL-1 (red) and DAPI (blue). Merged images are shown in the right panel. **Labelling**: A, amygdala; BV, blood vessel; Cr, crypts, also termed glands; Ctx, cerebral cortex; d3V, dorsal third ventricle; Hi, hippocampus; HN, habenular nuclei; HT, hypothalamus; IE, intestinal epithelium LV, lateral ventricle; LP, lamina propria; MM, muscularis mucosae; Th, thalamus; v3V, ventral third ventricle; Vi, villi.

**Extended Data Fig 8.**
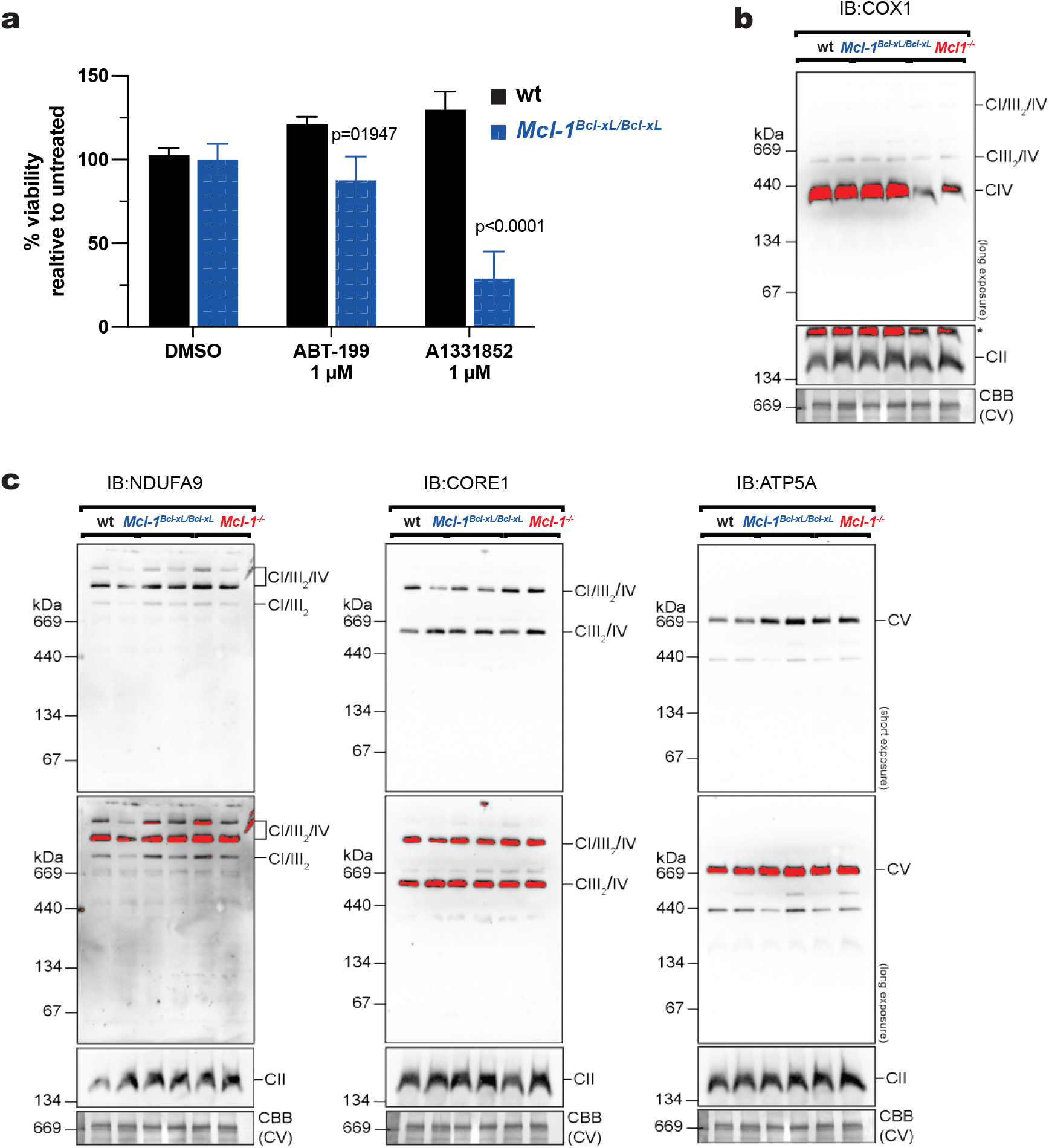
BCL-XL can replace the function of MCL-1 in the assembly of the electron transport chain. **a**, Survival analysis of immortalised wild-type and homozygous *Mcl-1^Bcl-xL/Bcl-xL^* MEFs (n=2 per genotype). Cell viability was assessed 48 h after treatment with the indicated BH3 mimetic drugs by MTT assay. Data are presented as mean ±SEM from 2 independent experiments performed in triplicates. Statistical significance was tested using two-way ANOVA with Tukeys’s multiple comparison. p-values are indicated in the figure. **b**, Blue Native Page and immunoblot analysis for respiratory complex IV (CIV) (using an antibody to COX1) on isolated mitochondria from wild-type (wt), *Mcl-1^-/-^* and *Mcl-1^Bcl-xL/Bcl-xL^* MEFs. A portion of the Coomassie stained blot (CBB) with complex V (CV) shown serves as a loading control. This is a long exposure of the same blot presented in Fig. 5C. **c**, Blue Native Page and immunoblot analysis for respiratory complexes (using antibodies to NDUFA9, CORE1, ATP5A or SDHA (SDHA shown in lower box labelled CII)) on isolated mitochondria from wild-type, *Mcl-1^-/-^* and *Mcl-1^Bcl-xL/Bcl-xL^* MEFs, with both short and long exposures. A portion of the Coomassie stained blot (CBB) with complex V (CV) shown serves as a loading control.

**Extended Data Movies 1-3** Representative movies of wild-type **(Movie 1),** homozygous *Mcl-1^Bcl-xL/Bcl-xL^* **(Movie 2)** and homozygous *Mcl-1^Bcl-2/Bcl-2^* **(Movie 3**) E12.5 embryos (C57BL/6 background) after whole-mount embryo staining and clearing (iDISCO^39^). The vasculature is marked by CD31/PCAM1 staining (red) and blood cells are marked in green due to their inherent autofluorescence.

## Extended Data Tables

**Extended Data Table 1:** E11.5-E14.5 Anomalies.

|  |  | *normal* | *Small, normal* | *Small, abnormal* | *Small, dead* | *cranial abnormalities other than exencephaly* | *exencephaly* | *blood accumulation* |
| --- | --- | --- | --- | --- | --- | --- | --- | --- |
| E11.5 | *wild type* | 34 (37) | 2 (37) | 1 (37) | 0 | 0 | 0 | 1 (37) |
|  | *Mcl-1<sup>Bcl-xL/Bcl-xL</sup>* | 3 (8) | 1 (8) | 1 (9) | 1 (8) | <u>2 (9)</u> | 0 | 2 (9) |
|  | *Mcl-1<sup>Bcl-2/Bcl-2</sup>* | 4 (17) | 0 | 4 (17) | 7 (17) | 0 | 2 (17) | 5 (17) |
| E12.5 | *wild type* | 36 (36) | 0 | 0 | 0 | 0 | 0 | 0 |
|  | *Mcl-1<sup>Bcl-xL/Bcl-xL</sup>* | 10 (10) | 0 | 0 | 0 | 0 | 0 | 0 |
|  | *Mcl-1<sup>Bcl-2/Bcl-2</sup>* | 7 (12) | 0 | 0 | 0 | 0 | 1 (7) | 4 (12) |
| E13.5 | *wild type* | 12 (12) | 0 | 0 | 0 | 0 | 0 | 0 |
|  | *Mcl-1<sup>Bcl-xL/Bcl-xL</sup>* | 6 (8) | 0 | 0 | 0 | 0 | 0 | 2 (8) |
|  | *Mcl-1<sup>Bcl-2/Bcl-2</sup>* | 4 (4) | 0 | 0 | 0 | 0 | 0 | 0 |
| E14.5 | *wild type* | 9 (12) | 0 | 0 | 2 (12) | 0 | 0 | 1 (12) |
|  | *Mcl-1<sup>Bcl-xL/Bcl-xL</sup>* | 3 (3) | 0 | 0 |  | 0 | 0 | 0 |
|  | *Mcl-1<sup>Bcl-2/Bcl-2</sup>* | 3 (5) | 0 | 0 | 1 (5) | 0 | 0 | 2 (5) |

**Extended Data Table 2:**
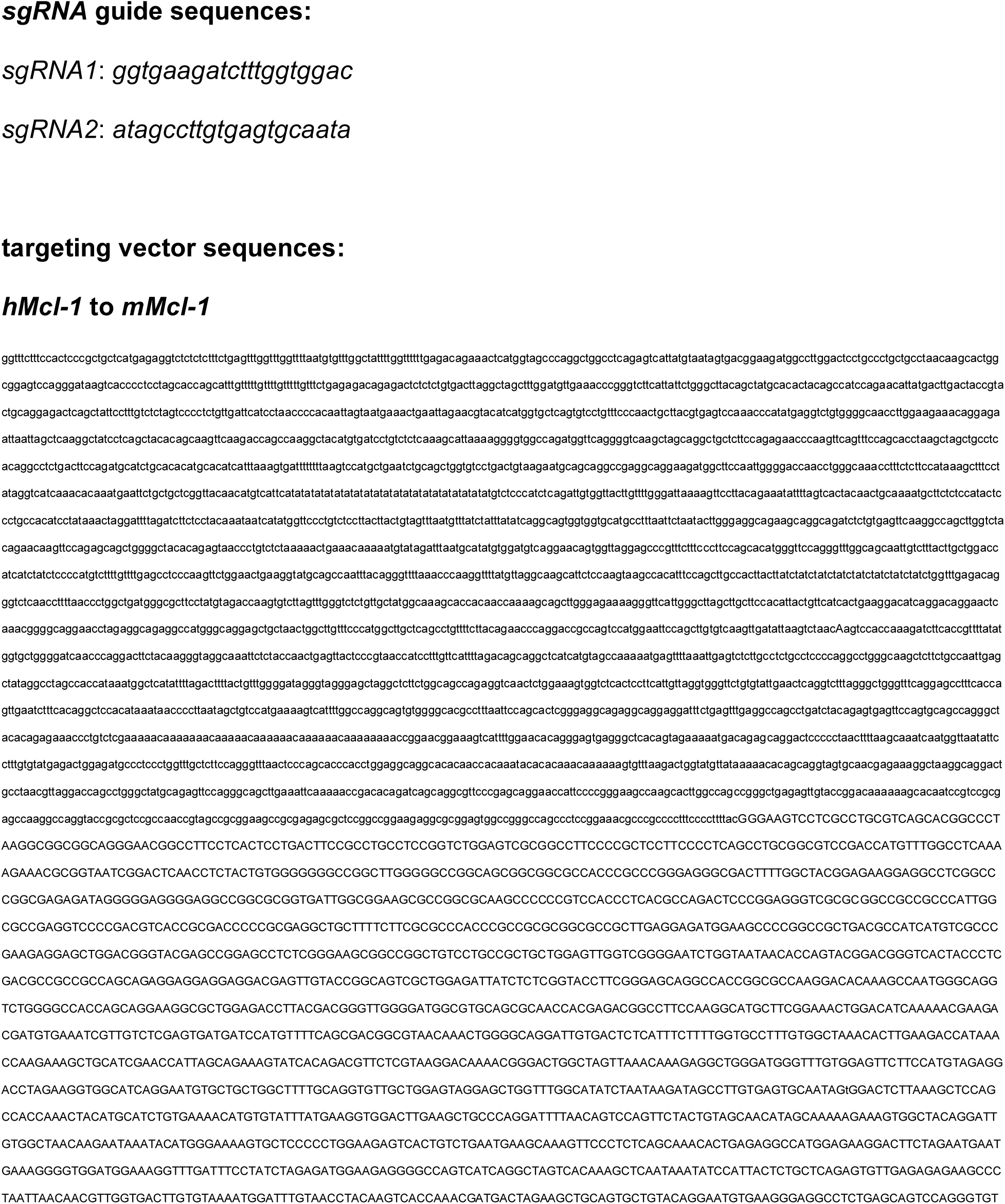
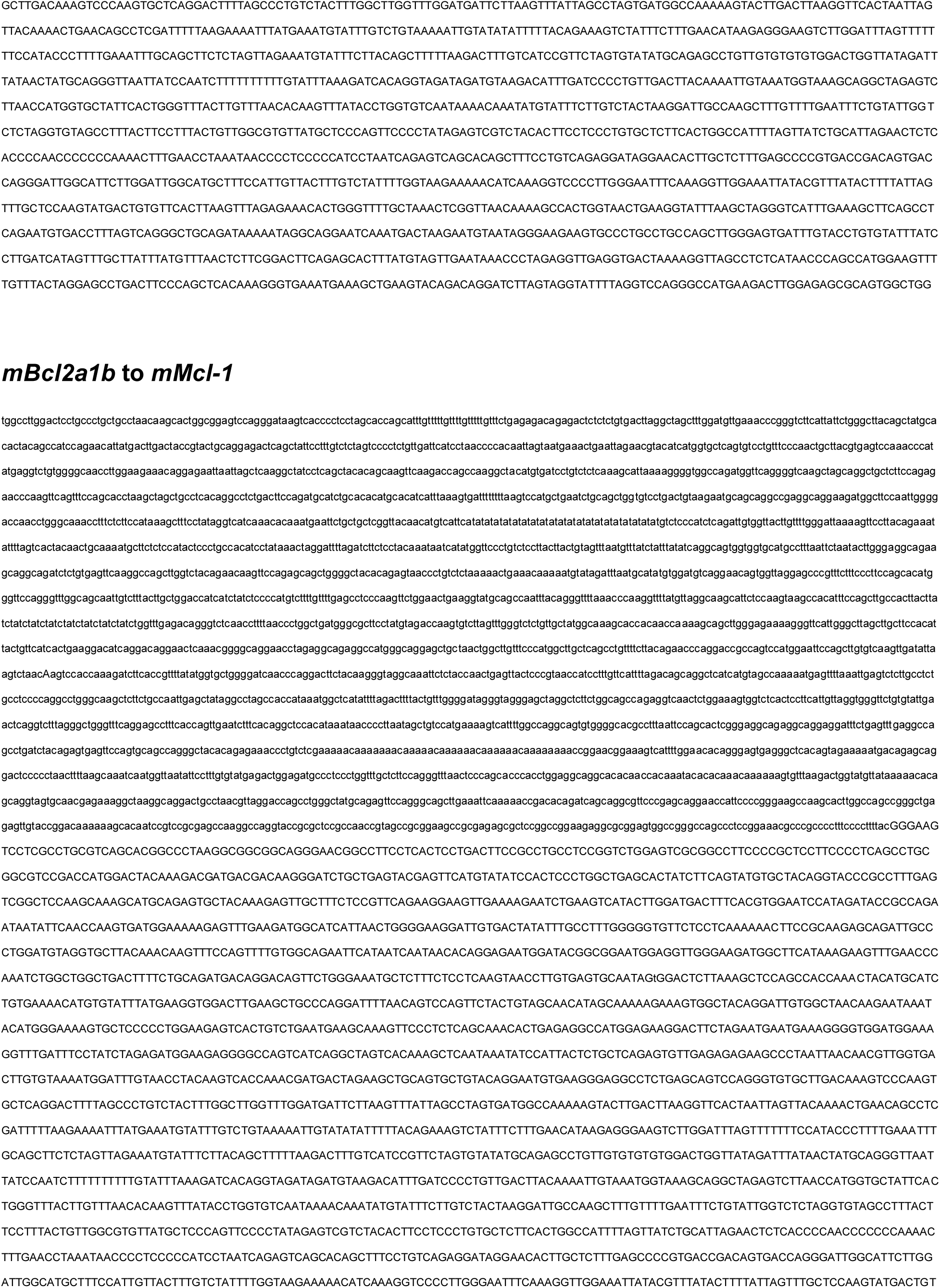
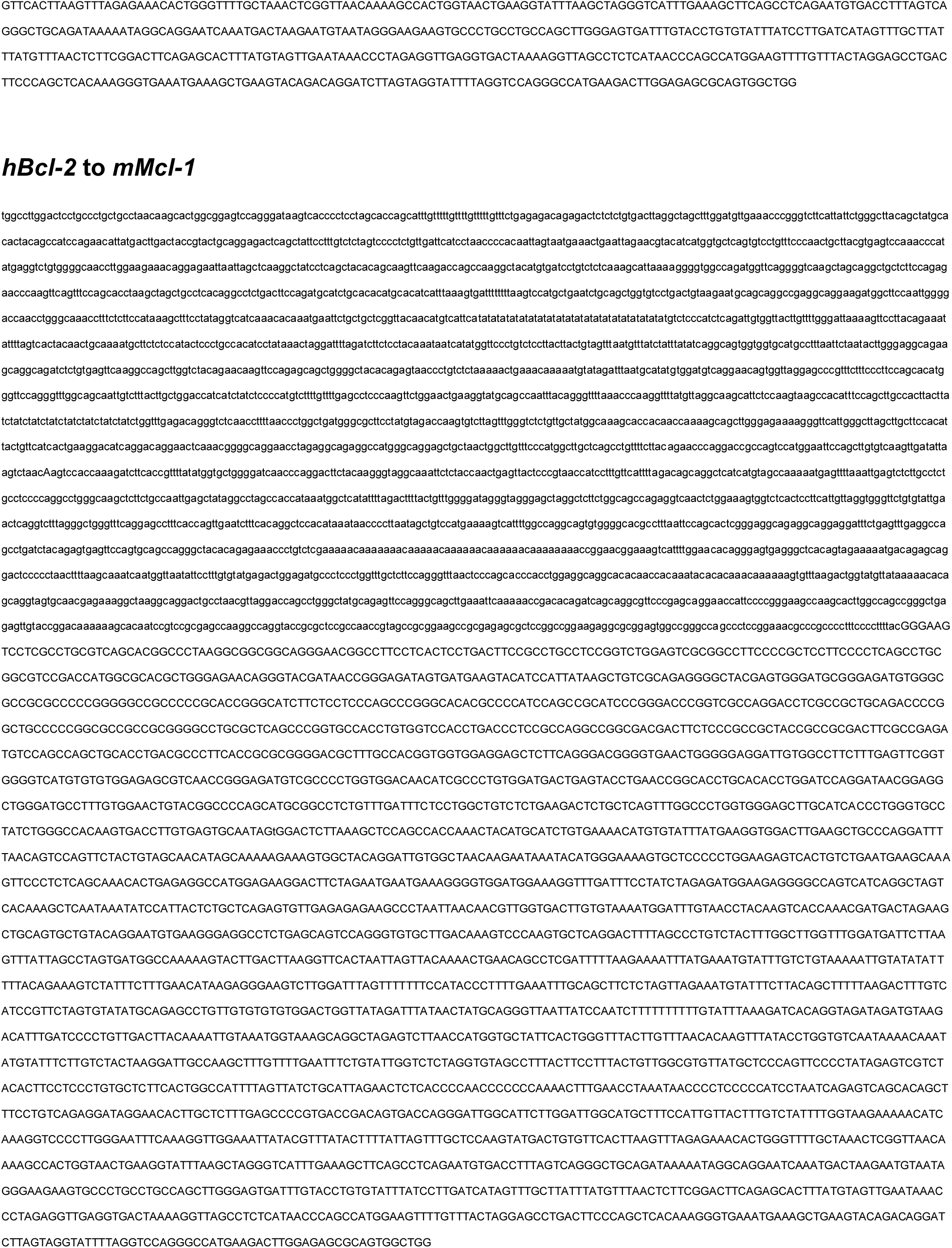
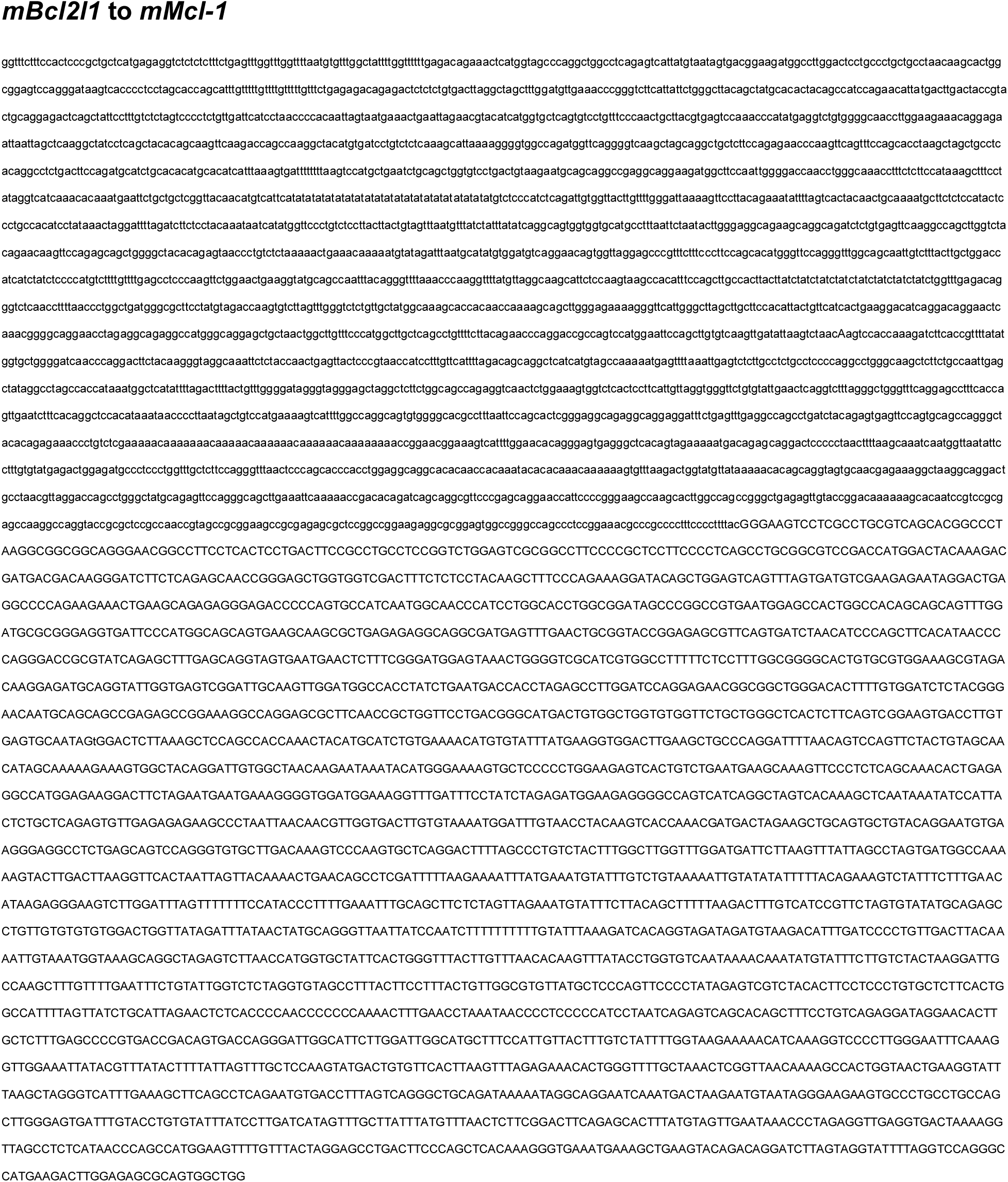
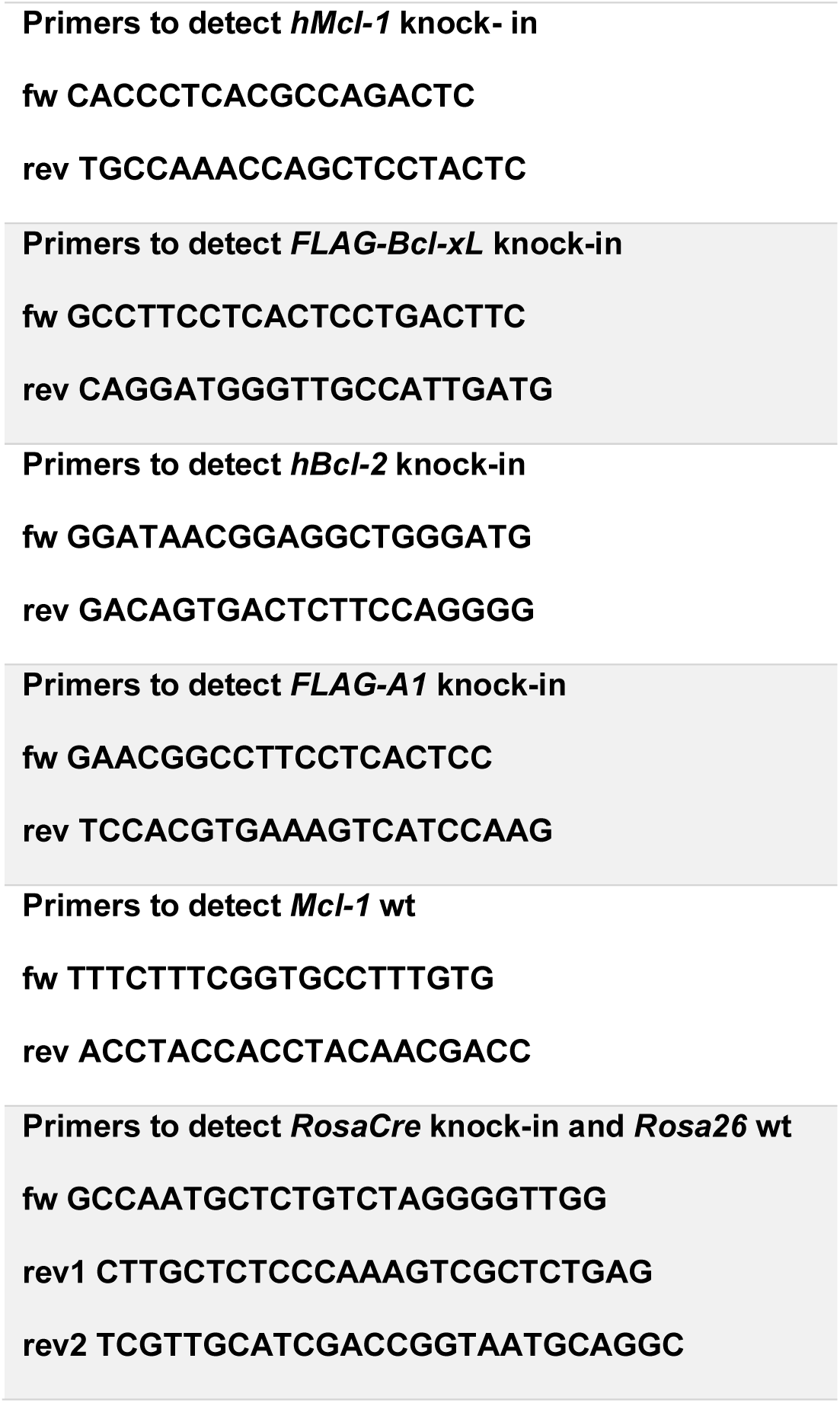
*sgRNA* guide, targeting vector and oligonucleotide sequences used for genotyping.

**Extended Data Table 3:**
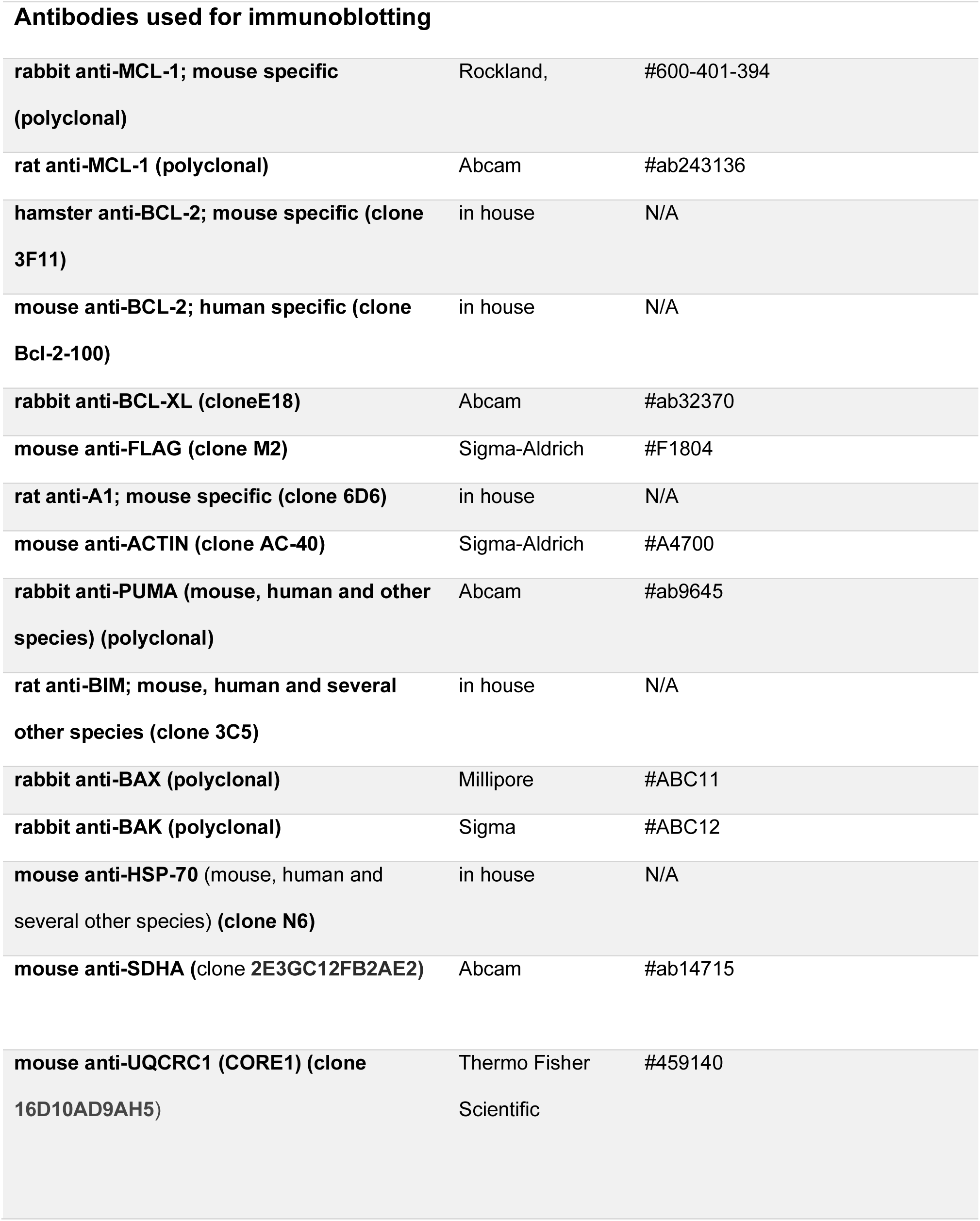
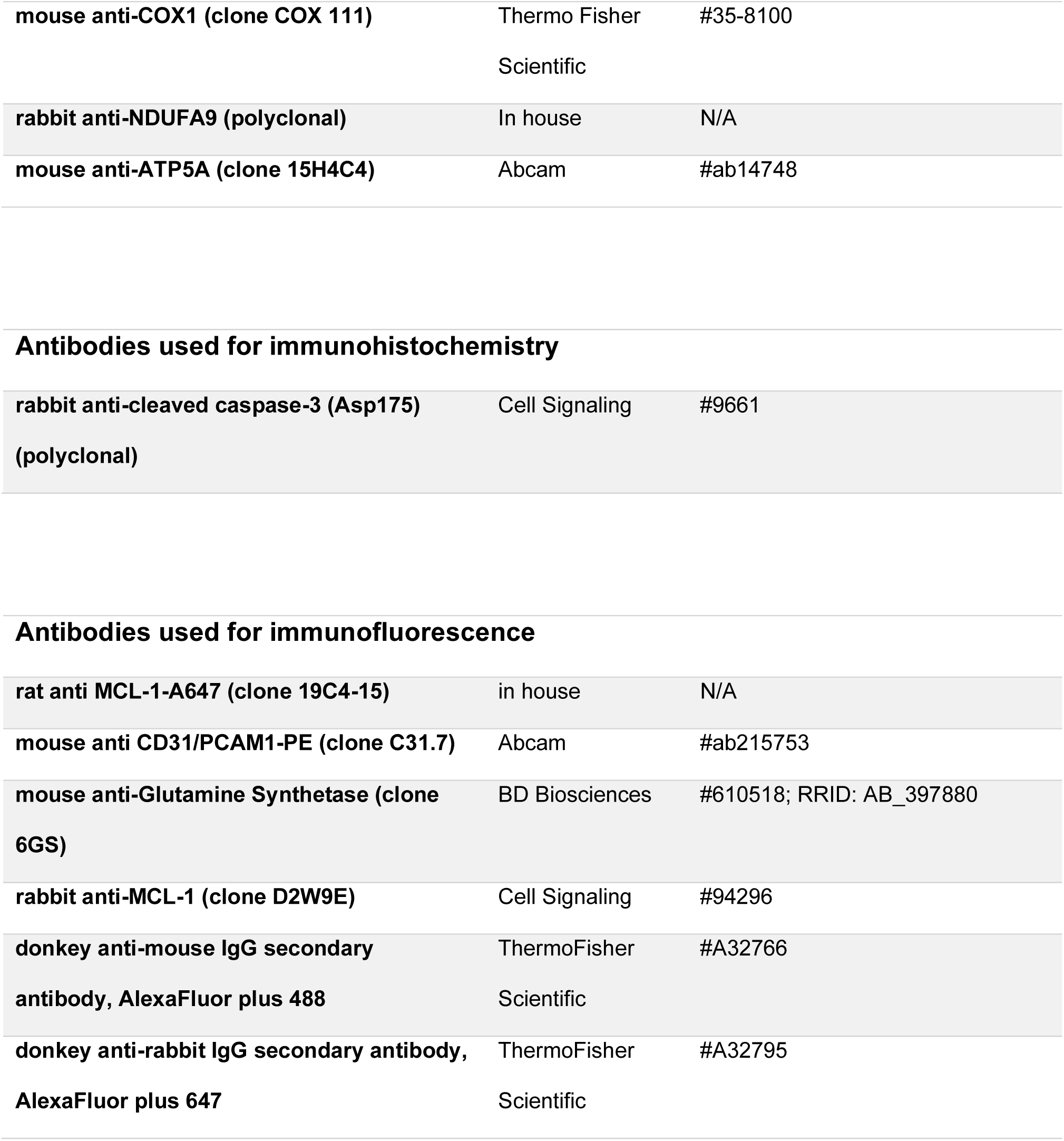
List of primary antibodies.

| **Antibodies used for immunoblotting** |  |  |
| --- | --- | --- |
| **rabbit anti-MCL-1; mouse specific (polyclonal)** | Rockland, | #600-401-394 |
| **rat anti-MCL-1 (polyclonal)** | Abcam | #ab243136 |
| **hamster anti-BCL-2; mouse specific (clone 3F11)** | in house | N/A |
| **mouse anti-BCL-2; human specific (clone Bcl-2-100)** | in house | N/A |
| **rabbit anti-BCL-XL (clone E18)** | Abcam | #ab32370 |
| **mouse anti-FLAG (clone M2)** | Sigma-Aldrich | #F1804 |
| **rat anti-A1; mouse specific (clone 6D6)** | in house | N/A |
| **mouse anti-ACTIN (clone AC-40)** | Sigma-Aldrich | #A4700 |
| **rabbit anti-PUMA (mouse, human and other species) (polyclonal)** | Abcam | #ab9645 |
| **rat anti-BIM; mouse, human and several other species (clone 3C5)** | in house | N/A |
| **rabbit anti-BAX (polyclonal)** | Millipore | #ABC11 |
| **rabbit anti-BAK (polyclonal)** | Sigma | #ABC12 |
| **mouse anti-HSP-70 (mouse, human and several other species) (clone N6)** | in house | N/A |
| **mouse anti-SDHA (clone 2E3GC12FB2AE2)** | Abcam | #ab14715 |
| **mouse anti-UQCRC1 (CORE1) (clone 16D10AD9AH5)** | Thermo Fisher Scientific | #459140 |
| **mouse anti-COX1 (clone COX 111)** | Thermo Fisher Scientific | #35-8100 |
| **rabbit anti-NDUFA9 (polyclonal)** | In house | N/A |
| **mouse anti-ATP5A (clone 15H4C4)** | Abcam | #ab14748 |
| Antibodies used for immunohistochemistry |  |  |
| **rabbit anti-cleaved caspase-3 (Asp175)**<br>**(polyclonal)** | Cell Signaling | #9661 |
| Antibodies used for immunofluorescence |  |  |
| **rat anti MCL-1-A647 (clone 19C4-15)** | in house | N/A |
| **mouse anti CD31/PCAM1-PE (clone C31.7)** | Abcam | #ab215753 |
| **mouse anti-Glutamine Synthetase (clone 6GS)** | BD Biosciences | #610518; RRID: AB_397880 |
| **rabbit anti-MCL-1 (clone D2W9E)** | Cell Signaling | #94296 |
| **donkey anti-mouse IgG secondary antibody, AlexaFluor plus 488** | ThermoFisher Scientific | #A32766 |
| **donkey anti-rabbit IgG secondary antibody, AlexaFluor plus 647** | ThermoFisher Scientific | #A32795 |

